# Inferring diffusion, reaction, and exchange parameters from imperfect FRAP

**DOI:** 10.1101/2025.05.06.652329

**Authors:** Enrico Lorenzetti, Celia Municio-Diaz, Nicolas Minc, Arezki Boudaoud, Antoine Fruleux

## Abstract

Fluorescence recovery after photobleaching (FRAP) is broadly used to investigate the dynamics of molecules in cells and tissues, notably to quantify diffusion coefficients. FRAP is based on the spatiotemporal imaging of fluorescent molecules following an initial bleaching of fluorescence in a region of the sample. Although a large number of methods have been developed to infer kinetic parameters from experiments, it is still a challenge to fully characterize molecular dynamics from noisy experiments in which diffusion is coupled to other molecular processes or in which the initial bleaching profile is not perfectly controlled. To address this challenge, we have developed HiFRAP to quantify the reaction-(or exchange-)diffusion kinetic parameters from FRAP under imperfect experimental conditions. HiFRAP is based on a low-rank approximation of a kernel related to the model Green’s function and is implemented as an ImageJ/Python macro for (potentially curved) one-dimensional systems and for two-dimensional systems. To the best of our knowledge, HiFRAP offers features that have not been combined together: making no assumption on the initial bleaching profile, which does not need to be known; accounting for the limitation of the optical setup by diffraction; inferring several kinetic parameters from a single experiment; providing errors on parameter estimation; and testing model goodness. In the future, our approach could be applied to other dynamical processes described by linear partial differential equations, which could be useful beyond FRAP, in experiments where the concentration fields are monitored over space and time.

**SIGNIFICANCE:** Fluorescence recovery after photobleaching (FRAP) is a microscopy approach that is widely used to investigate the diffusion and transport of molecules in life sciences and in material sciences. Numerous methods have been developed to derive kinetic parameters such as diffusion and binding coefficients. However, these methods suffer from limitations associated with experimental constraints, such as technical noise or an imperfectly known initial condition. To circumvent these limitations, we developed a comprehensive approach to estimate several kinetic parameters from a single experiment, to assess the precision of estimation, and to test whether the underlying model is well-suited. We implemented this approach in HiFRAP, an ImageJ/Python macro of broad applicability to one- and two-dimensional systems.

## 1 INTRODUCTION

Cells and tissues are the place of permanent transport and transformation of matter. At cellular level, trafficking, binding and unbinding, or diffusion, are essential in the self-organization of the cell, for instance. At multicellular level, diffusion, directed transport, and degradation of morphogens are key to setting morphogen distributions and providing positional information during organism development. Several methods have been developed to assess such molecular dynamics, including fluorescence recovery after photobleaching (FRAP), fluorescence spectroscopy, or single-particle tracking (1). Among these, FRAP appears as the most widely used method (1–5), likely because the microscopy setup is technically accessible.

FRAP is designed to study the dynamics of fluorescent molecules by monitoring the response to an initial perturbation. Molecules are first photobleached by strong and short light pulses in a region of the sample, often circle- or square-shaped. This causes a drip in light intensity re-emitted by the sample in this region. Fluorescence is then followed over space and time by time-lapse imaging with a microscope. Typically, the fluorescence (partially) recovers its initial level, and the pattern of recovery is informative of the underlying dynamics. When molecules only undergo diffusion, the timescale of fluorescence recovery is a function of the diffusion coefficient and of the size and shape of the initially bleached region (we will often use FRAPped region in the following) (3–5). Accordingly, FRAP is routinely used to determine diffusion coefficients. When molecules only undergo binding and unbinding to immobile substrates, then the timescale of fluorescence recovery is the inverse of the binding rate (3–5). Here, we consider more complex situations where several molecular processes are coupled, such as diffusion and binding/unbinding.

At cellular level, FRAP has also been used to investigate protein synthesis (6), dynamics of molecular condensates (7), mechanosensing (8), transport of mRNA (9), or cell adhesion (10, 11). At the tissue level, FRAP has been used to assess diffusion of morphogens (12) or expansion of the extra-cellular matrix (13). FRAP also appears in material science, for instance to characterize pharmaceutical compounds (3, 14). Despite the practical importance of FRAP, a comprehensive method to analyze and interpret FRAP data is still lacking (3). Here, we contribute to tackling this issue.

The classical method to determine diffusion coefficient is based on the theoretical calculation of the average concentration in the bleached region and fits to the recovery curve of fluorescence in that region (15, 16). Although the classical method is easy to implement, it assumes that the initial bleaching profile is perfectly known, it does not use the information available in spatial variations, and it does not easily allow to distinguish diffusion from other processes (15). These limitations prompted the development of more sophisticated methods. Methods that do not require the knowledge of the bleaching profile are based on the decomposition of the fluorescence levels into Fourier modes and fit the temporal decay of the mode amplitude to theoretical solutions, in linear (17, 18), in axisymmetric geometry (19), or without assumptions on geometry (20). Methods that use all spatial information to improve precision use fits of the spatiotemporal concentration field to analytical (21) or numerical (22, 23) solutions of the diffusion equation, the former being restricted to Heaviside-like initial bleaching profile and the latter allowing initial bleaching profile of arbitrary shape. Other studies built methods to account for the effect of boundary conditions on diffusion (7), for diffusion on curved surfaces (24, 25), or for anomalous diffusion (26, 27).

Several studies have addressed the use of FRAP to determine the kinetic parameters of chemical reactions, binding/unbinding dynamics, or exchanges between compartments, which all formally amount to chemical reactions. In general, these studies directly deduce constants from average pre-bleaching fluorescence and recovery time of average fluorescence in the bleached area (6, 28–30), possibly accounting for rapid diffusion before the reactions take place. However, there are discrepancies between values of kinetic parameters according to the model used (31) and it is difficult to disentangle reactions from diffusion (32).

Another line of investigation has accounted for the coupling between reactions and diffusion. It is possible to solve numerically reaction-diffusion equations in complex realistic geometries to simulate FRAP and investigate changes in qualitative behavior according to parameters (33). To obtain kinetic parameters, an option is to use fluorescence recovery in the bleached area or in the region of interest and to fit analytical recovery curves of several experiments with varying sizes of the bleached regions, which provides enough information to deduce more than one kinetic parameter (34–36). Another option is to use all spatial information and fit the spatiotemporal concentration field (fluorescence level) to analytical (26, 37) or numerical (38–41) solutions of reaction-diffusion equations (for one or two species, according to the problem of interest). We note that these methods are constrained by the need to know the initial condition, i.e. the profile of fluorescence following initial bleaching. Using Fourier coefficients of the fluorescence field (42) like in some of the methods to infer diffusivity already mentioned, it was possible to get rid of this constraint, at the price of averaging several experiments together to average out noise. We aim at going beyond limitations by noise and the need to precisely know the bleached profile. Indeed, the bleached profile is difficult to control experimentally (20) and discrepancies with the assumed profile generally lead to a misestimation of kinetic parameters (16, 36).

Some studies accounted for other couplings, such as advection (directed transport) and diffusion (15), advection-reaction (43), or advection-reaction-diffusion (9), with similar limitations to those previously discussed. Here we only consider reaction-diffusion, but we note that our method is generalizable to any process described by linear partial differential equations. In addition, we account for photobleaching during imaging, i.e. bleaching of fluorophores due to their excitation during time-lapse imaging, following a few studies that inferred the rate of photobleaching during imaging from experimental data (19, 22, 40). Altogether, we aim at building a method to infer from FRAP the kinetic parameters of a process described by a reaction-diffusion equation from single experiments. We assume that experiments are noisy, that the initial bleaching profile is unknown, and that gradual photobleaching occurs due to imaging. We also account for the diffraction-limited resolution of the optical setup. In the following, we formulate the problem as one of cost minimization and propose a systematic method to solve it. We validate and optimize the approach with synthetic data. Finally, we present the implementation of the approach as an ImageJ macro (available on https://github.com/lorenzetti1996/HiFRAP_project) and illustrate it with experimental data on the the transmembrane protein Mtl2p in fission yeast (44).

## 2 RESULTS AND DISCUSSION

### 2.1 General framework in the case of pure diffusion

FRAP experiments involve imaging at regular intervals a region of interest containing the initially bleached (FRAPped) domain. The quantity of light emitted from the sample is proportional to the local concentration of fluorescent molecules, as long as saturation of the detectors is avoided. However, imaging the sample also results in photobleaching, so that fluorescence is attenuated by a constant factor after each image. The optical setup causes a spatial smoothing of the light pattern due to diffraction, which is characterized by the point-spread function of the microscope. The recorded signal results from three contributions. The first contribution is proportional to the concentration of fluorescent molecules, provided appropriate tuning of the excitation laser and detector gain; it contains a noisy part due to the statistical fluctuations of the number of these molecules in a small volume. The second contribution originates in technical noise, mostly associated with the detector. The third contribution is a background homogeneous signal associated with the detector. Accordingly, the recorded signal is a smoothed version of the field of fluorophore concentration, combined with noise and shifted by a background intensity. In HiFRAP, we account for point-spread function, photobleaching during imaging due to imaging, background intensity, and noise. The time-lapse data are assumed to be a temporal sequence of square images. The signal to be analyzed is sampled *n*_*t*_ times after every time interval Δ*t*, over *n*_*x*_ × *n*_*x*_ square pixels of side length Δ*x*.

For the sake of simplicity, we introduce and validate HiFRAP assuming that the underlying dynamics is set by diffusion only. We generalize our approach to reaction-diffusion in Section 2.6. We illustrate our approach in two dimensions, although the ImageJ/Python plugin is also implemented for diffusion along a line. The concentration of fluorophores *c*(*x, y, t*) is a function of spatial coordinates (*x, y*) and of time *t*. It is a solution of the diffusion equation

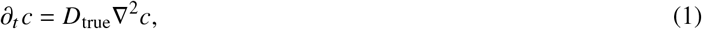

where *D*_true_ is the true diffusion coefficient, *∂*_*t*_ is the partial derivative with respect to time and ∇^2^ the Laplace operator. Our goal is to provide the best estimate *D*_est_ of the diffusion coefficient.

When photobleaching during imaging is not negligible, homogeneous regions of the sample (far from the FRAPped domain) show fluorescence decaying by a factor *ρ*_*i*_ = exp (−*εt*_*i*_ /Δ*t*), where *ε* is the decay rate per image, *t*_*i*_ the time at which the *i*-th image is collected, and Δ*t* is the time interval between two consecutive images. Accordingly, the theoretical solution of the diffusion equation (1), *c* (*x, y, t*_*i*_) should be multiplied by this factor *ρ*_*i*_. Photobleaching, however, does not affect the background intensity, *I*_BG_, which is considered to be constant in time and space. The background value *I*_BG_ and the decay rate per image *ε* are either supposed to be known from regions distinct from the FRAPed area as explained in Section 2.7, or inferred altogether with the dynamical parameters as discussed in Section 2.6.

Figure 1**A** shows an example of the synthetic dataset generated by solving analytically the diffusion equation (1); concentration is spatially smoothed to mimic the effect of diffraction and account for the point-spread function, see Section 4.8 for details. The region of interest is a square of side length *L*. The FRAPped domain is a square of side length *ℓ* = *L* / 3 in which the fluorophores concentration is set to 0 at *t* = 0. The first row shows the simulated spatial profile of the fluorophore concentration, which becomes smoother and converges to the initial density over time, as could be expected. The second row shows a microscope-like time-lapse imaging, obtained from the simulated (true) concentration field by adding an uncorrelated Gaussian random variable corresponding to technical noise and applying a Gaussian spatial filtering corresponding to the point-spread function of the optical setup, see Section 4.8. In the following, we use such synthetic data to test our method and estimate the precision of the estimated diffusion coefficient, *D*_est_, with respect to its true value, *D*_true_.

**Figure 1:**
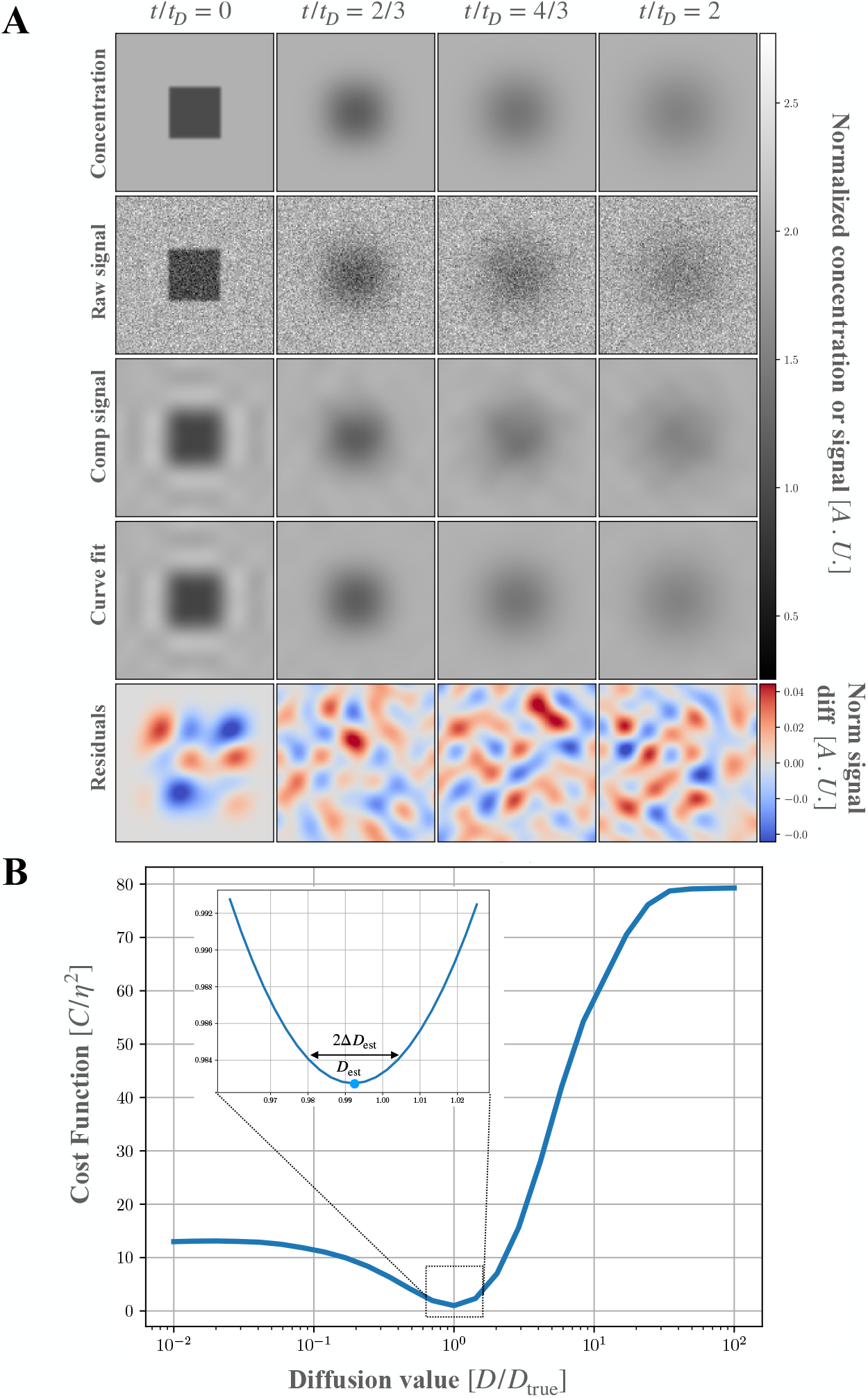
Inferring the diffusion coefficient from simulated FRAP. **A**. A square region of interest of side length *L* is monitored and a central square region of side length *ℓ* = *L* / 3 is FRAPped at *t* = 0. From top to bottom: raw artificial data, microscope-like synthetic data accounting for diffraction and technical noise, compressed synthetic data, concentration field fitted by HiFRAP, and residuals of the fit. From left to right: snapshots from *t* = 0 to *t* = 2*t*_*D*_, where 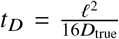 is taken as a unit of time. Gray- and color-scales indicate the concentration or signal intensity normalised by the drop in concentration Δ*I* at *t* = 0 in the FRAPped square. Dark (blue) to bright (red) indicate low to high concentration or signal. The decay rate per image due to photobleaching is set to *ε* = 0. For other parameters, default values are given in Section 4.8. **B**. Cost function *C* (normalised by noise amplitude *η*) as a function of fitting diffusion coefficient *D* (normalized by its true value *D*_true_), with a magnification of the neighbourhood of the minimum of *C* in the inset. The cost function is minimal at *D*_est_, which is close to *D*_true_, up to an estimated error Δ*D*_est_. The total observation time *T* for this dataset is *T* = 2*t*_*D*_ and the time step Δ*t* = 0.133*t*_*D*_.

### 2.2 A method to infer parameters from FRAP experiments

HiFRAP estimates kinetic parameters such as diffusion coefficient independently of any assumptions on the initial bleaching pattern. We use *N*_tot_ = *n*_*x*_ × *n*_*x*_ × *n*_*t*_ (2D spatial × temporal) pixels from time-lapse imaging. We fit a theoretical model to those data by minimising a cost function that quantifies the differences between observed data and theoretical solution. The model is built from the solution for (1) and accounts for the point-spread function (see Section 2.1 for more details).

The fit of the model to the data is performed iteratively in two steps, each minimising the cost function: at each iteration, we first estimate the initial conditions for fixed kinetic parameters, and then update the kinetic parameters (in this case, the diffusion coefficient). To estimate the initial conditions, we use both spatial and temporal data and define the cost function as the sum of squared deviations between model predictions and the observed data. Because the partial differential equation is linear with respect to the initial condition, the model response is also linear, and the cost function becomes quadratic in the initial condition. As a result, for fixed kinetic parameters, the optimal initial condition can be obtained by solving a linear system of equations. Once the cost is minimized with respect to the initial conditions (for fixed kinetic parameters), it is normalized by the effective number of degrees of freedom (see Section 4.2.4). The kinetic parameters are then updated using the reflective trust region algorithm, a least-squares optimization method, to minimize the normalized cost.

Two important subtleties arise in the estimation of initial conditions at each iteration. First, the problem is inherently ill-posed: an infinite number of initial conditions can produce the same observed data, since measurements are finite while the initial condition is a continuous field belonging to an infinite-dimensional space. To address this, we use pseudo-inversion to identify the components of the initial condition that influence the observed data. This reveals that only a limited number of degrees of freedom in the initial pattern significantly affect the measurements. Second, pseudo-inversion becomes computationally expensive for large datasets, as it involves operations on matrices of size *N*_tot_ × *N*_tot_, which is prohibitive when *N*_tot_ *>* 10^5^. To overcome this, we reduce the data size through compression, enabling implementation in an interactive ImageJ plugin. Specifically, we apply a discrete Fourier transform to compress the data in both spatial dimensions and retain only a subset of Fourier components. This reduces the dataset to a manageable size *N* = *n*_*q*_ ×*n*_*q*_ × *n*_*t*_ . As shown in Section 2.4, we find that values of *n*_*q*_ between 5 and 9 are sufficient to obtain reliable estimates of the kinetic parameters. The resulting compressed dataset, with *N* on the order of 10^3^ to 10^4^, allows efficient pseudo-inversion to compute the optimal initial condition for the current estimate of the kinetic parameters (here the diffusion coefficient *D*).

To assess the uncertainty of the inferred parameter, we estimate the curvature of the cost function around the minimum by computing the Hessian as a higher curvature indicates higher precision (Figure 1B), see Section 4.2.5 for details. To evaluate the goodness of the fit we use two metrics. The adjusted coefficient of determination 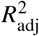 is the classical coefficient of determination (*R*^2^) corrected using the effective number of degrees of freedom; it quantifies how well the model explains variations of the observed data, and ranges from 0 (no explanation) to 1 (perfect fit). A limitation of the 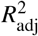 is that its value may depend on the amplitude of noise and on the initial bleaching profile (see section 4.2.5). Accordingly, the 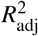 should be used to compare two models applied to the same experiment, as used in section 2.6 and 2.7. The second metrics is not sensitive to the level of noise and the bleaching profile as it estimates whether the distribution of residuals is compatible with a Gaussian distribution using the Kolmogorov-Smirnov test. It returns a p-value indicating whether to reject the null hypothesis that the residual distribution is compatible with a Gaussian. The only limitation is that it is computationally very slow because it involves an explicit calculation of all residuals in an appropriate basis, see Section 4.2.5.

HiFRAP is illustrated with synthetic data in Figure 1. In panel **A**, the two first rows are the true concentration of fluorophores and the simulated microscope-like images. For our analysis, we compress the microscope-like images (third row) to which we fit the model (fourth row), obtaining relatively small residuals (fifth row). In panel **B**, we plot the cost function (after optimizing the initial condition) as a function of the fitting parameter *D* for the diffusion coefficient. The estimated diffusion coefficient *D*_est_ is defined as the value of *D* that minimizes the cost. We use the curvature of the cost function (inset in panel **B**) and noise amplitude to estimate the uncertainty Δ*D*_est_ on the diffusion coefficient, see Section 4.2.5. Here, the coefficient of diffusion is well estimated, *D*_est_ /*D*_true_ = 0.992, and the relative uncertainty Δ*D*_est_ /*D*_est_ = 0.01 is small. Finally, we test the validity of the model by examining whether residuals are normally distributed, which is implemented using the Kolmogorov-Smirnov tests (45), see Section 4.2.5. Here we find a p-value of 0.57, showing that diffusion model is a good model for these data, as could be expected. Moreover, the adjusted coefficient of determination has a value 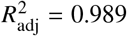, though it is only meaningful when comparing two models applied to a single dataset.

HiFRAP was also implemented for FRAP experiment along a line. In Figure S6, we show an example of HiFRAP inference applied to sythetic data for a 1D diffusive system, with similar results to the 2D case.

### 2.3 Validation on artificial data

To thoroughly evaluate the robustness of the inference method, we applied HiFRAP to six collections of synthetic datasets, each corresponding to a different noise level. Within each collection, multiple datasets were generated using identical model parameters (as in Figure 1) but with different realizations of the noise. For each synthetic dataset, we estimated the diffusion coefficient *D*_est_ and its error Δ*D*_est_. For each collection, we computed the average estimate ⟨*D*_est_⟩ (the brackets ⟨ ⟩ stand for average over the collection) and the empirical error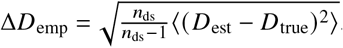

In Figure 2, we plotted the estimated diffusion coefficient *D*_est_ and estimated uncertainty Δ*D*_est_ as a function of noise strength. Panel **A** shows that the distribution of *D*_est_ is well centered around its true value *D*_true_. The standard deviation of this distribution increases with the noise amplitude *η* and the coefficient of variation Δ*D*_emp_/*D*_est_ reaches values comparable to 1 for noise strengths *η* such that 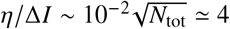, where Δ*I* is the amplitude of the drop in intensity following initial photobleaching. Panel **B** shows that the estimated error Δ*D*_est_ agrees well with the empirical error Δ*D*_emp_, except for the highest amplitude of noise, validating the estimation of error from a single dataset. HiFRAP also works for diffusion along a line as shown with synthetic data in Figure S6, although with less precision due to less spatial averaging (less pixels) than in 2D diffusion. Overall, HiFRAP provides good estimates of the diffusion coefficient and of its error from a single experiment.

**Figure 2:**
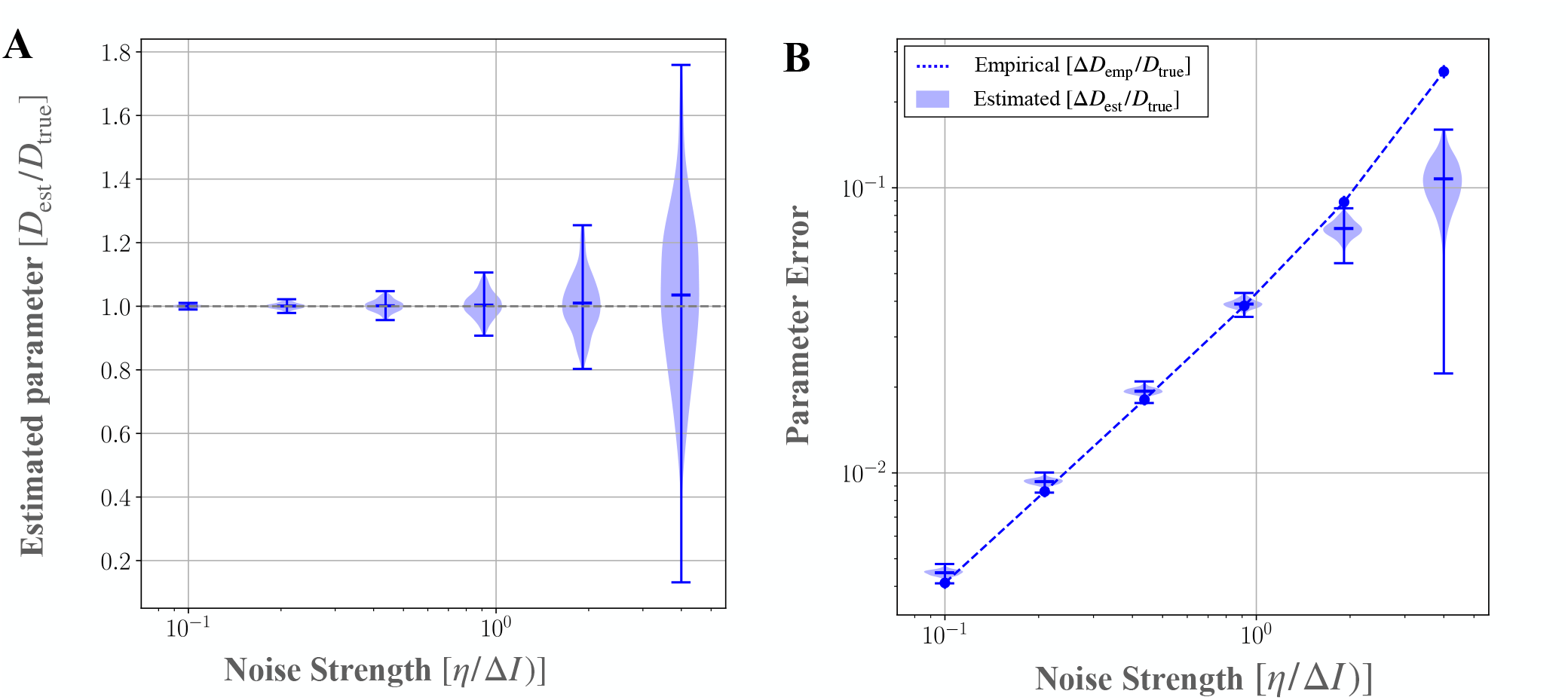
Validation of HiFRAP on a collection of synthetic data. **A** Estimated diffusion coefficient *D*_est_; **B** estimated uncertainty Δ*D*_est_ and empirical error Δ*D*_emp_. The quantities are all normalised by the true diffusion constant *D*_true_ and plotted as a function of the normalised noise amplitude *η* / Δ*I*, where *η* is noise amplitude, Δ*I* is the drop-off in intensity after initial photobleaching. Violins represent distributions of *D*_est_ and Δ*D*_est_ while the ticks highlight average and extreme values. The dashed gray line in **A** represents the reference value *D*_est_ /*D*_true_ = 1, while the dashed blue line in **B** corresponds to the empirical error Δ*D*_emp_. The number of realisations is 200 for each value of noise strength and the decay rate per image due to photobleaching is set to *ε* = 0.

### 2.4 Optimizing experimental and analysis parameters

To optimize the estimates of kinetic parameters (here diffusion coefficient), we aim at tuning parameters that are accessible in experiments — side length of FRAPed region *ℓ*, number of time frames *n*_*t*_, and delay between frames Δ*t* — and in analysis — number of modes *n*_*q*_ kept for the compression. We implicitly assume the space resolution, Δ*x*, to be constrained by the microscope used and the side length of the region of interest, *L*, to be constrained by the size of the system and its spatial variations — the region of interest should be as big as possible while sufficiently homogeneous. The bleaching size *ℓ* and the number of modes *n*_*q*_ influence the amount of useful spatial information. To be optimal, the size of the FRAPped region *ℓ* should be large enough for the perturbation in fluorescence associated with FRAP to be of significant weight compared to noise, whereas *ℓ* comparable to *L* leads to a loss of spatial information contained in the periphery of the FRAPped region. We found that the error on estimation of the diffusion coefficient is minimal roughly around *ℓ* ∼ *L*/3 in the case of a square FRAPped region. Concerning the number of modes kept in the discrete Fourier transform, fair estimates are reached for *n*_*q*_ ≥ 5 (Figure S2). Accordingly, we took *ℓ* = *L*/3 and *n*_*q*_ = 9 in all our analyses, except when specified otherwise.

The time resolution of experiments may be constrained by the sample imaged, for instance when there is phototoxicity. Here we only consider constraints due to the optical setup, which are mostly associated with photobleaching during imaging. The intensity of the observed signal decays with the number of images acquired, proportionally to exp [−*ε* (*n*_*t*_ −1)], so that the signal quickly vanishes when *n*_*t*_ increases beyond 1 /*ε* + 1. We therefore choose *n*_*t*_ to be the integer part of 1 /*ε* + 1. Concerning the choice of the time step Δ*t* between two images, we note that the temporal decay rate due to photobleaching is *ε*/ Δ*t*, while the relaxation (to equilibrium) rate due to diffusion is the inverse of the diffusion time *t*_*D*_ = *ℓ*^2^ /(16*D*) — defined as the time at which the standard deviation of the position of a Brownian particle reaches half the side length of the FRAPed region. If the decay due to photobleaching is high, then fluorescence disappears before diffusion can be observed, whereas if the decay rate due to photobleaching is low, most of the images are taken after diffusion has homogenised concentrations and these images are not informative. As a consequence, we expect the optimal delay between images to correspond to *ε* / Δ*t* 1 ∼ /*t*_*D*_. To further test this conclusion, we plotted in Figure 3 the normalized empirical error Δ*D*_emp_ /*D*_emp_ as function of dimensionless delay Δ*t* /(*t*_*D*_*ε*) . As expected, the plot shows that the error Δ*D*_emp_ has a minimum. This minimum occurs when Δ*t* ∼10*t*_*D*_*ε*, a value that we used in the remainder of this study.

**Figure 3:**
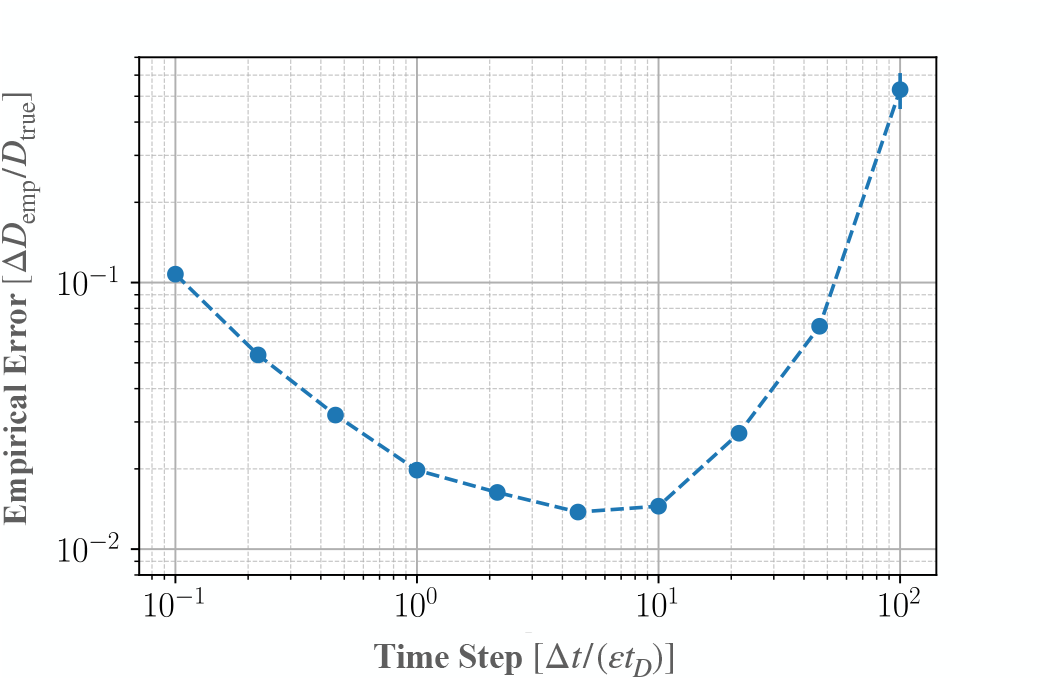
Optimization of imaging time step Δ*t*. The empirical error Δ*D*_emp_, normalised by the true diffusion coefficient *D*_true_, is represented as a function of relaxation time Δ*t*/*ε* due to photobleaching during imaging, normalised by the diffusive timescale *t*_*D*_ = *ℓ* ^2^/*D*_true_/16. The empirical error was computed as 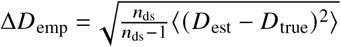 from the estimated values *D*_est_ and true value *D*_true_ of the diffusion coefficient from *n*_ds_ = 200 artificial datasets.

### 2.5 Benchmarking HiFRAP

To test the efficiency of HiFRAP, we compared it to classical methods to obtain diffusion coefficient. Beforehand, we stress the versatility of HiFRAP because it makes no assumption on FRAP patterns nor on boundary conditions, which is not the case of classical approaches. This is illustrated in Figure S3, where the region FRAPped is axisymmetric with a Gaussian profile, X-shaped, or E-shaped. For the comparison with other methods, which assume the bleaching pattern to be known, we considered a square FRAPped domain. Benchmarking was performed with respect to two classical approaches, as described in (16, 36). The two approaches are based on determining the bleaching profile by fitting the post-bleach image, and then estimating the diffusion coefficient by analysing the recovery curve, i.e. the temporal variations of the signal average over the FRAPped region. For the first step, we followed (36) and modelled the square FRAPped profile as a 2D sharp function *f* (*x, y*) = *H* (∓ /2 −*x* + *x*_0_) *H* (*x* −*x*_0_ + ∓ / 2) *H* (∓/ 2 −*y* +*y*_0_) *H* (*y* −*y*_0_ + ∓ /2) (with *H* the Heaviside function *H* (*u*) = 0 if *u <* 0 and *H* (*u*) = 1 if *u >* 0), smoothed to account for the point-spread function of the microscope. Accordingly the post-bleach profile is parametrised by its size *ℓ*, the position of its center (*x*_0_, *y*_0_) and the smoothing length. In a second step the two approaches differ. In the relaxation time based method, the half-recovery time *τ*_1_ / _2_ is estimated as the time at which the intensity has recovered half of its initial value in the FRAPped region (16). The diffusion coefficient is then computed using to the theoretical relation 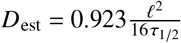 valid for a square FRAPped region. In the temporal fit based method, the diffusion coefficient is obtained by fitting the recovery curve to the theoretical curve for a FRAPped region with side length and smoothing length, as obtained in the first step. The final formula is found in (36).

As the three methods retrieve the true diffusion coefficient on average, we compared their respective precisions. The empirical errors on diffusion coefficient are shown in Figure 4 for different amplitudes of noise. The errors are computed for 200 artificial datasets for each noise strength. As could be expected, the empirical error increases with the noise strength. HiFRAP performs better than the two other approaches, especially at lower noise values, although HiFRAP does not make any assumption on the FRAPped bleaching profile. We ascribe the performance of HiFRAP to the fitting of spatiotemporal data instead of temporal data in other approaches.

**Figure 4:**
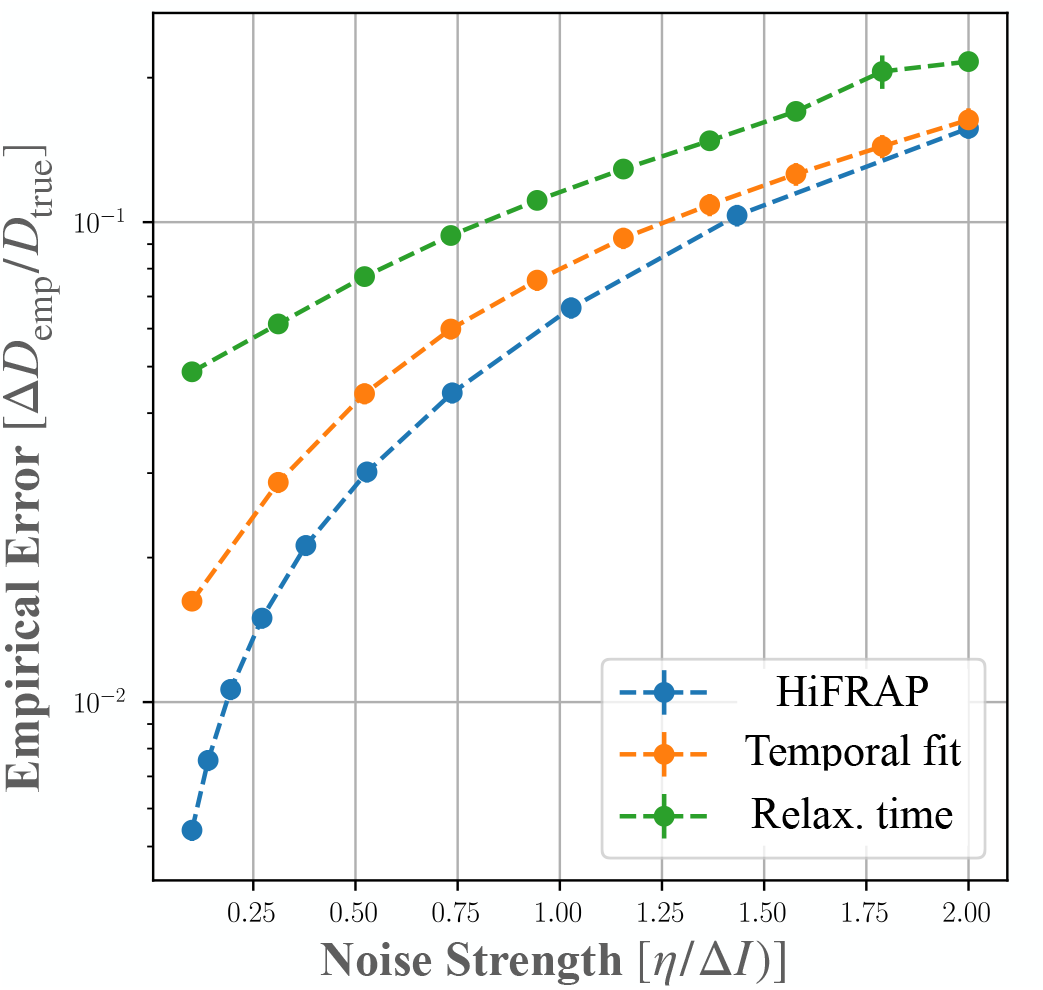
Benchmarking of HiFRAP. Empirical error Δ*D*_emp_ as function of the noise strength *η*, the noise amplitude, normalised in terms of signal drop upon bleaching Δ*I*. The lines and relaxation-time-based (green), temporal-fit-based (orange), and HiFRAP (blue) methods. Empirical error was computed as 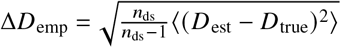 from estimated values *D*_est_ *D*_true_ and true value of the diffusion coefficient from *n*_ds_ = 200 number of datasets.

### 2.6 Applying HiFRAP to reaction-diffusion

Our method can be generalized to infer kinetic parameters for more complex dynamics. Besides diffusion, molecules may be synthesised, degraded, or undergo other chemical reactions. In addition, membrane-localised proteins or lipids may be exocytosed or endocytosed. As long as the changes in concentration are not too large, the dynamics of one chemical species can be modelled by a linear diffusion-reaction equation,

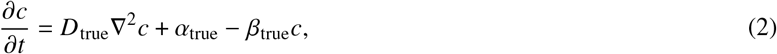

where *D*_true_ is still the diffusion coefficient, while *α*_true_ and *β*_true_ represent source rate and exchange rate, respectively, both assumed to be constant. The interpretation of these reaction terms depends on context. For instance, *α*_true_ may correspond to a synthesis rate and *β*_true_ to a degradation rate. In the case of a membrane-localised molecule, *α*_true_ and *β*_true_ may correspond to the rates of exocytosis and endocytosis, respectively, of this molecule. To extend HiFRAP to reaction-diffusion, we followed the same approach as in Section 2.2 with the difference that the cost to minimize now depends on multiple parameters, *D, α, β*, and/or *ε*. The minimisation with respect to *D, β* and *ε* is performed numerically using the reflective trust region algorithm, while *α* and *I*_BG_ is computed analytically, see Section 4.5 for details. The imaging step can be optimised as in Section 2.4. Equation (2) involves two characteristic times, the diffusion time, *t*_*D*_ = *ℓ*^2^ /(16*D*_true_), and the exchange time *t*_*β*_ = log (2) / *β*. The typical recovery time *t*_*r*_ for the combined dynamics is expected to be of the order of 1/ *t*_*r*_ = 1/ *t*_*D*_ + 1/ *t*_*β*_. Accordingly, the optimal time step can be taken as Δ*t* ∼*εt*_*r*_ /10.

We generated artificial reaction-diffusion data with the same initial square FRAP profile as before, as described in Section 4.8. Figure 5 shows the inference of the parameters of Eq. (2) from these data using HiFRAP. We first assumed that the stationary solution *c*_s_ is perfectly known *a priori*, from the pre-bleaching images for instance. When *c*_s_ is known, the cost function depends only on *α* and *β*, see Section 4.5. In panel **A**, we show an example of the contours of the cost function. The estimate on *α* may be obtained from the relation *α*_est_ = *c*_s_ *β*_est_. To explore the behavior of our method, we would *a priori* need to vary *D*_true_, *α*_true_ and *β*_true_. However changing the ratio *α*_true_ /*β*_true_ only changes the stationary concentration and so does not affect the uncertainty of the different estimates. Changing the recovery time *t*_*r*_ does not significantly affect the precision of the method, since we adapt the time resolution of the experiment accordingly. We therefore varied the ratio *t*_*β*_ /*t*_*D*_ in Figure 5B-C, keeping *α*_true_ /*β*_true_ and *t*_*r*_ constant. As could be expected (37), the uncertainty on the diffusion coefficient becomes high when the fluorophore dynamics is dominated by reaction and, reciprocally, the uncertainty on the relaxation coefficient becomes high when the dynamics is dominated by diffusion. We note that, like in the pure diffusion case, the curvature of the cost function yields a good estimate of the error on parameters, see Figure (S1). Finally, we assessed the quality of the fit by comparing it to fits to diffusion-only (assuming *α*_true_ = *β*_true_=0) and reaction-only models (assuming *D*_true_ = 0) using the adjusted coefficient of determination 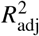. Figure 5D shows that 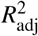 is always much larger for the reaction-diffusion model, meaning that it better explains the data, except for low or high ratios of reaction to diffusion times, for which the reaction-diffusion model is almost equivalent to one of the two simplest models.

**Figure 5:**
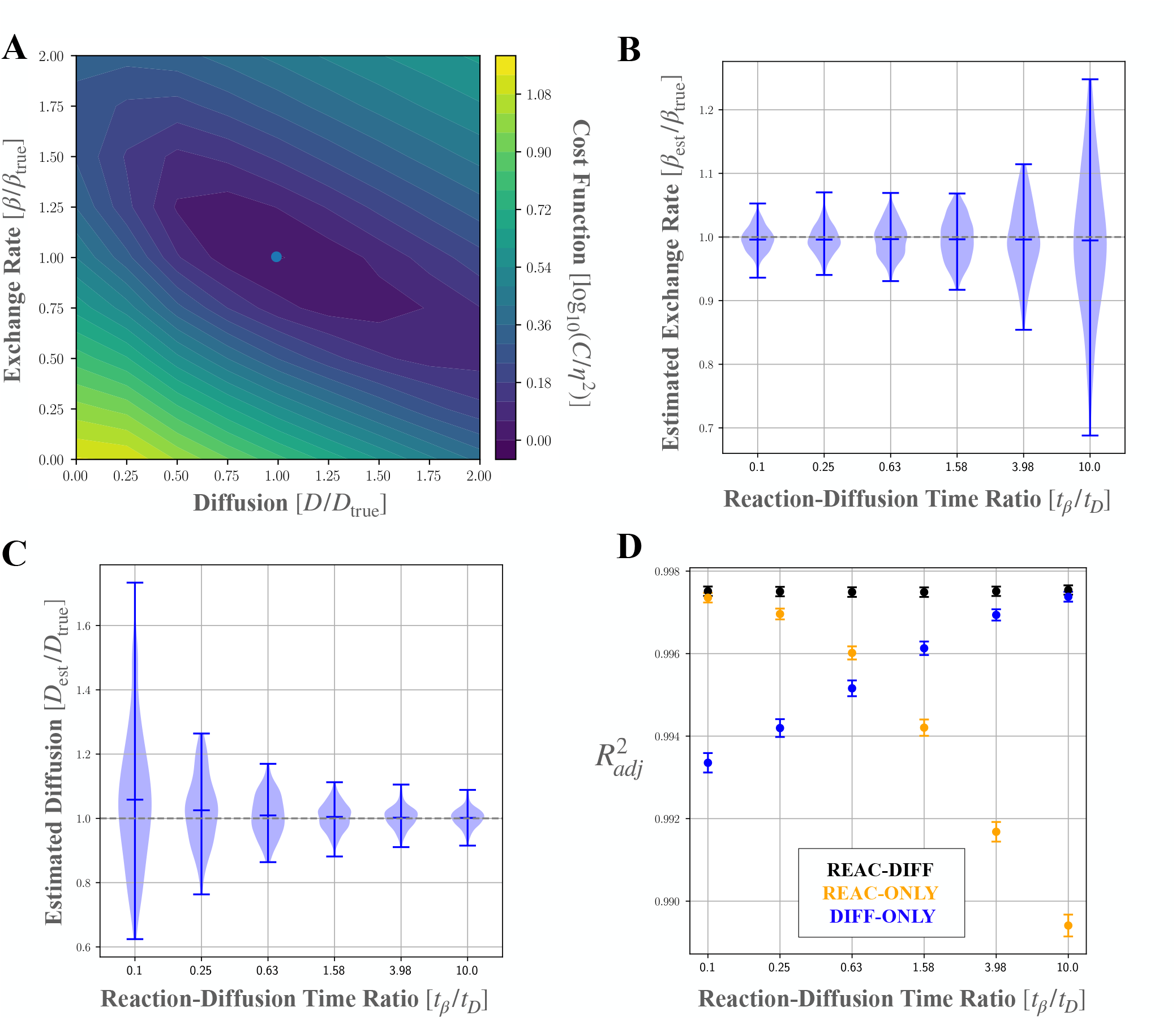
HiFRAP applied to the estimation of diffusion and reaction rates knowing the pre-FRAP concentration field and the rate of photobleaching during imaging. **A** Contour plot of the normalized cost function (log_10_ [*C* /*η*^2^], with *η* the amplitude of the noise) as a function of the normalized fitting parameters *D* /*D*_true_ and *β* /*β*_true_, while the source term *α* is constrained to be *α* = *βc*_*s*_, with *c*_*s*_ the stationary concentration known from prebleaching images. The colorscale is shown on the right, with blue and yellow corresponding to low and high cost, respectively. The light blue point indicates the minimum of the cost function; its coordinates yield the estimates *D*_est_ and *β*_est_. **B-C** Estimated dynamical parameters *D*_*est*_ and *β*_*est*_ as function of the reaction-diffusion time ratio *t*_*β*_ /*t*_*D*_ = 16 log (2) /*ℓ*^2^*D*_true_ /*β*_true_ keeping the signal relaxation time *t*_*r*_ = (1 /*t*_*β*_ +1 /*t*_*D*_)^−1^ = *ε*Δ*t* / 7 constant. Violin plots show the distribution of the estimates and horizontal ticks indicate the maximum, the average, and the minimum of the distribution. The dashed gray line represents the reference value at which the estimated parameter is equal to the true parameter of the system. Here, the stationary concentration *c*_*s*_ is assumed to be known from the average pre-FRAP concentration field, so that *α*_est_ = *c*_*s*_ *β*_est_. The rate of photobleaching during imaging, *ε* is supposed to be known from a control area. The number of datasets analysed is 200. **D** Adjusted coefficient of determination, 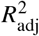, for three different fits, diffusion only, reaction only and reaction-diffusion, as a function of reaction-diffusion time ratio. The error bar represents the mean and the standard deviation of the distribution.

When the values of rate of photobleaching during imaging, stationary concentration and background are unknown, they can be estimated together with the kinetic parameters using optimisation in 5-dimensional space. Figure 6 shows the results of HiFRAP in this case. Our estimates remain fairly good. The errors on diffusion and reaction coefficients behave like in the preceding case, though they are bigger here owing to the highest dimensionality (5 instead of 2). The errors on the source rate, bleaching rate and background increase when the fluorophore dynamics is dominated by diffusion, similar to the exchange rate because all three parameters effectively relate to reactions.

**Figure 6:**
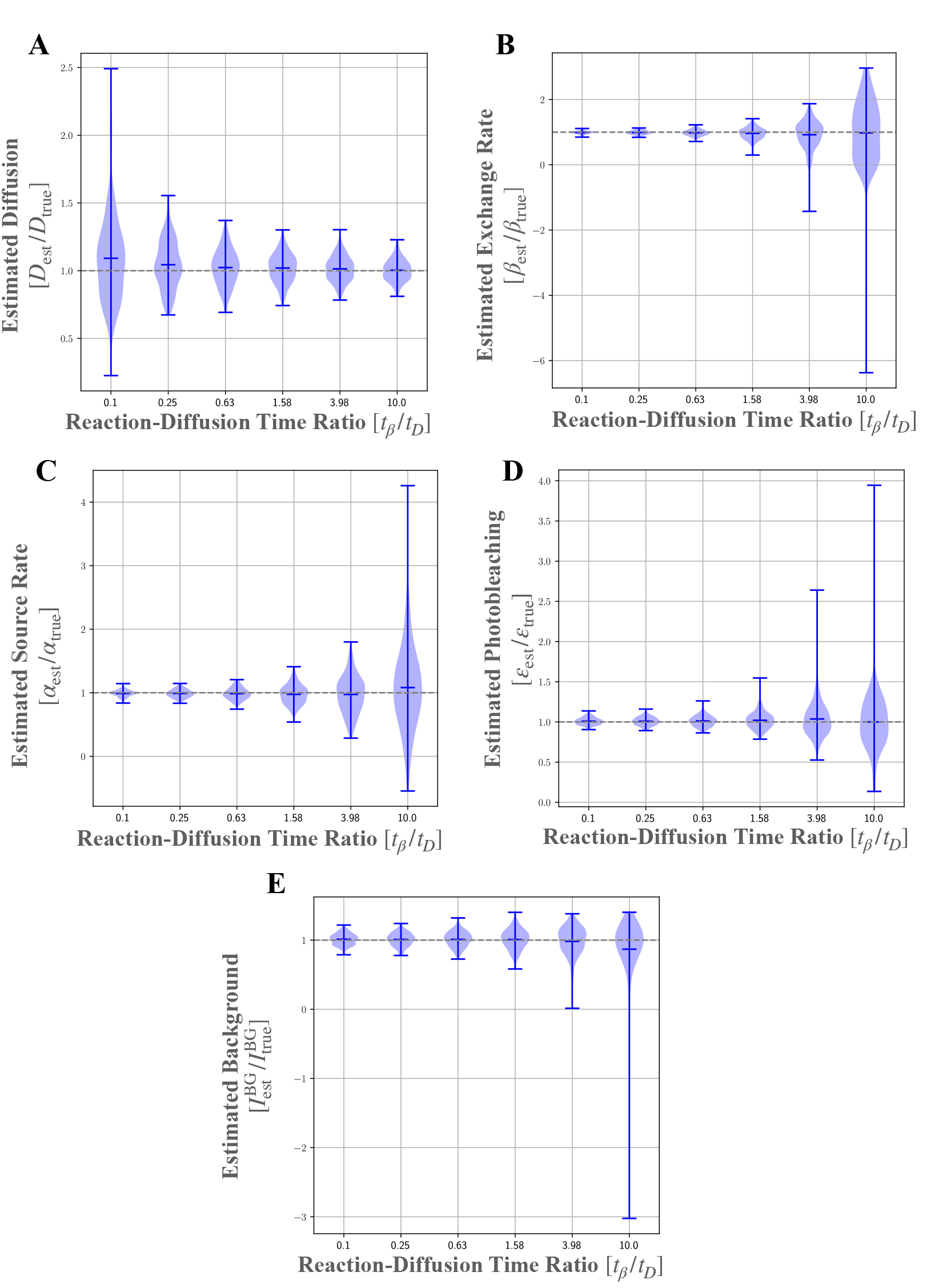
HiFRAP applied to diffusion-reaction when pre-FRAP concentration field and rate of photobleaching during imaging are unknown. Distributions of the estimated parameters (normalized by their true values) as a function of the reaction-diffusion time ratio *t*_*β*_ /*t*_*D*_ for 200 artificial datasets: **A** diffusion coefficient *D*_est_, **B** exchange rate *β*_*est*_, **C** source rate *α*_est_, and **D** rate of photobleaching during imaging *ε* and **E** background signal intensity *I*_BG_. Violin plots show the distributions. The horizontal ticks stand for the maximum, the average and the minimum of the distributions. The dashed gray line represents the reference value at which the estimated parameters is equal to the true value of the system.

### 2.7 Applying HiFRAP to experimental data

We implemented HiFRAP as an ImageJ macro that wraps Python scripts (available on https://github.com/lorenzetti1996/HiFRAP_project). Here we illustrate the macro with experiments in fission yeast. We considered a putative mechanosensitive transmembrane protein, Mtl2p, which is homogeneously distributed around the cell surface (44). We prepared and imaged cells as detailed in Section 4.9. Given that Mtl2p has very low cytoplasmic concentration, we hypothesised that, over the timescale of experiments, Mtl2p diffuses along the surface of the cell, and we aimed at testing this hypothesis and at estimating the diffusion coefficient.

Building upon preceding Sections, we implemented the HiFRAP ImageJ macro for three models (diffusion only, reaction-diffusion, or reaction only) and for various experimental options. These options include the potential use of additional information to reduce errors on the estimation of parameters, see Section 4.6 for details. In particular, HiFRAP enables to select two regions outside the region of interest (ROI — contains the FRAPped domain): a region free from fluorescent reporters (background ROI) — used to estimate the background value *I*_BG_ by averaging, and an unFRAPped region (control ROI) — used to estimate the rate of photobleaching during imaging *ε*. Figure 7A shows three example ROIs. The ROIs were chosen as large as possible to maximize the amount of information analyzed, within the constraint of being not too close to cell edges (as viewed from the top) or to image edges, to avoid the effects of, respectively, cell curvature and optical distortion on signal intensity. Potential additional information also includes prebleaching images, which yield stationary molecule concentration *c*_*s*_ by averaging.

**Figure 7:**
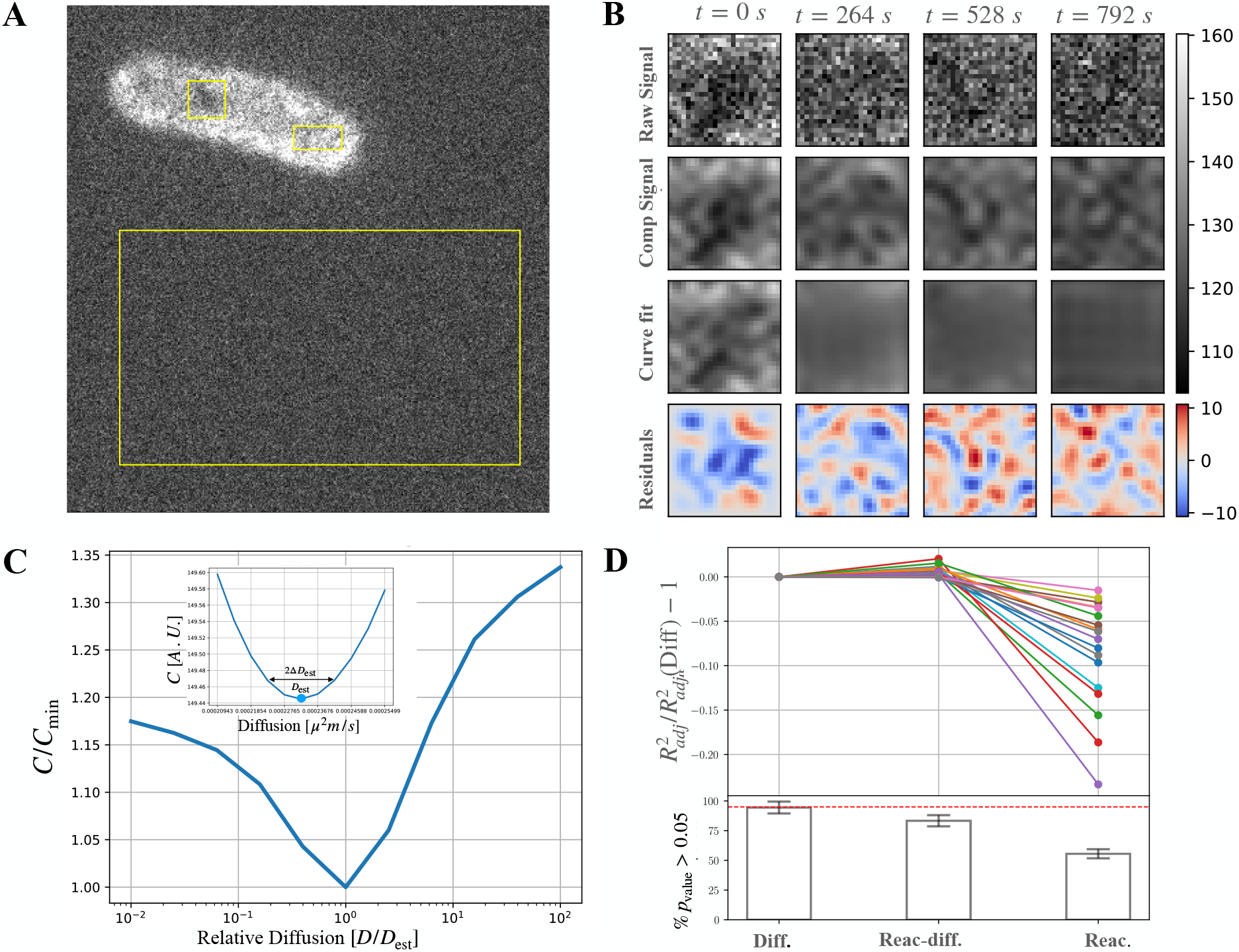
Inferring the diffusion coefficient from experimental data. **A** Fluorescence microscopy image of a fission yeast cell expressing Mlt2-GFP. The cell was bleached inside a region to the left of the cell centre. Three regions were selected with Fiji. The square is the region of interest that contains the FRAPped region and has side length *L* = 1.45 *µm*. The top rectangle is the control ROI, used to estimate rate of photobleaching during imaging. The bottom rectangle is the background ROI, used to estimate background signal intensity. **B** From experimental signal to fit; 2-dimensional data shown at 4 time points — time in seconds is indicated above the first row, *t* = 0 follows initial bleaching. From top to bottom: experimental data inside the ROI; compressed experimental data; fit to compressed experimental data; residuals of the fit. The grayscale and colorscale are shown on the right. **C** Corresponding cost function *C* (normalised by its minimum) as a function of fitting parameter *D* (normalised by the estimated diffusion coefficient *D*_est_) for the diffusion coefficient; the inset is a zoom-in around the minimum with the cost function in the arbitrary unit of the signal and the diffusion values in *µm*^2^ /*s*. **D** Assessing the goodness of the diffusion model by comparing it to reaction-diffusion and to reaction only models. Top: adjusted coefficient of determination 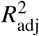 normalised using its value for the diffusion model; each line corresponds to one of the 18 experiments (one color per experiment). Kolmogorov test (bottom): each bar indicates the fraction of cases in which the Kolmogorov test was above the 0.05 value; the red dashed line indicates the level below which the model should be rejected; errorbars correspond the error in sampling a binomial distribution.

In FRAP experiments on Mtl2p-GFP, typical estimates are *I*_BG_ *≈* 100 *A*.*U*. for the background level (from the background ROI), *c*_*s*_ *≈* 30 [*A*.*U*.] for the stationary concentration (using one prebleaching image), and *ε* ∼10^−2^ for the photobleaching (using the control ROI). In Figure 7B-C we show the same plots as in Figure 1 for the estimation of the diffusion coefficient from one FRAP experiment. The cost function presents a single minimum (panel C) and the corresponding best fit to compressed experimental data is shown in the third row of panel B.

We optimized experimental parameters following Section 2.4. The size of the region of interest (ROI) being limited by cell width, we chose to use square ROIs of side length ∼ 1.5 *µm* and FRAPped square regions of side length *ℓ* ∼ 0.5 *µm*. In practice, we found that the FRAP was imperfect; the reduction in signal intensity was inhomogeneous and did not occur over a perfect square, see *t* = 0 in Figure 7B. This may be caused by different factors such as the small size of the FRAPped region, laser imprecision, or fluctuations in fluorophore concentration. To adjust the imaging time step Δ*t*, we first estimated the order of magnitude of the diffusion coefficient and of photobleaching rate per image by applying HiFRAP to a preliminary experiment. We found *ε* ∼10^−2^ and *D*_est_ ∼10^4^ *µm*^2^*s*^−1^. Then to optimize our analysis, we chose Δ*t* = (5 /8) *εℓ* ^2^ /*D* ∼ 10 *s*, and *n*_*t*_ = 1 /*ε* ≃ 100. Finally, to limit the computational cost, we used *n*_*q*_ = 9 modes for the compression.

Following parameter adjustment, each of the 18 analyzed cells was FRAPped once. We fitted experimental data to each of the three models: diffusion only, reaction diffusion, and reaction only. Estimated parameters are given in table 1. To assess whether the dynamics of Mtl2p-GFP is dominated by diffusion or by reaction, we computed the adjusted coefficient of determination 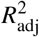, as shown in figure 7D. In all the 18 experiments, the diffusion model has a higher 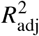 compared to the reaction model indicating that diffusion contributes more to Mtl2p dynamics at the experimental time-scale; the diffusion-reaction model has slightly higher 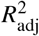 than the diffusion only model, suggesting that the two are almost equally good at explaining the data. Consistently, the Kolmogorov-Smirnov test rejects the reaction model (the p-value is greater than 0.05 for 60% of the cases, well below the threshold of 95%) and tends to reject the reaction-diffusion model. In addition, when considering a reaction-diffusion model, we find an average loss rate that is not positive, which is not physically acceptable because it means for instance that the rate of detachment of Mtl2p from the membrane is negative. For this reason, we conclude that pure diffusion better explains experimental data.

**Table 1:** Estimated parameters (mean ± standard deviation over 18 experiments) and associated errors for each model. From left to right: diffusion coefficient *D*_est_, error on diffusion 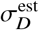, exchange rate *β*, error on loss 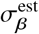, source rate *α*_est_ (AU stands for arbitrary unit), and error on source 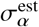. From top to bottom: reaction only, reaction-diffusion, and reaction only models.

| Parameter | $\langle D_{\text{est}} \rangle$ | $\langle \sigma_D^{\text{est}} \rangle$ | $\langle \beta_{\text{est}} \rangle$ | $\langle \sigma_\beta^{\text{est}} \rangle$ | $\langle \alpha_{\text{est}} \rangle$ | $\langle \sigma_\alpha^{\text{est}} \rangle$ |
| --- | --- | --- | --- | --- | --- | --- |
| Unit | $10^{-4} \mu\text{m}^2 / \text{s}$ | $10^{-4} \mu\text{m}^2 / \text{s}$ | $10^{-4} \text{s}^{-1}$ | $10^{-4} \text{s}^{-1}$ | AU $\text{s}^{-1}$ | AU $\text{s}^{-1}$ |
| Diff. | $1.93 \pm 1.40$ | $0.07 \pm 0.06$ | — | — | — | — |
| Reac.-Diff. | $3.3 \pm 3.2$ | $0.2 \pm 0.2$ | $-10 \pm 16$ | $2 \pm 1$ | $-0.18 \pm 0.29$ | $0.03 \pm 0.02$ |
| Reac. | — | — | $9.4 \pm 4.5$ | $0.5 \pm 0.2$ | $0.166 \pm 0.100$ | $0.008 \pm 0.003$ |

**Table 2:**
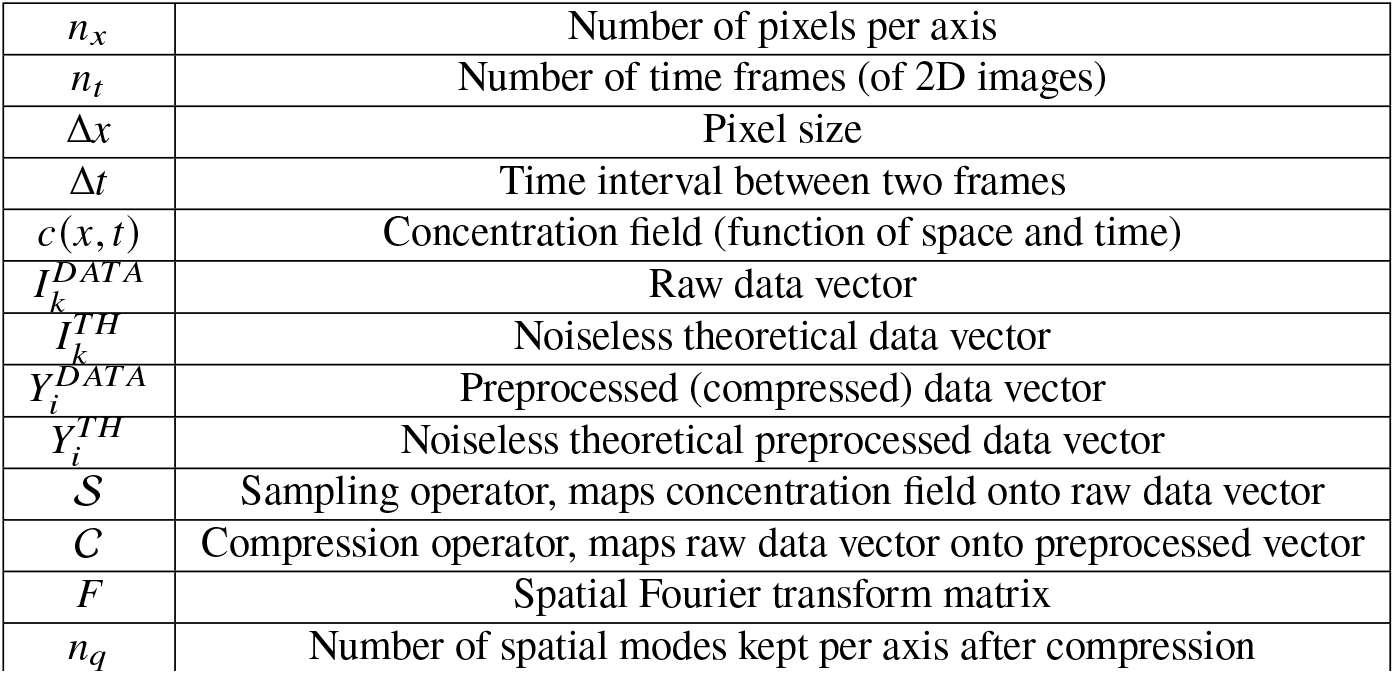
Notations for sampling and preprocessing.

|  |  |
| --- | --- |
| $n_x$ | Number of pixels per axis |
| $n_t$ | Number of time frames (of 2D images) |
| $\Delta x$ | Pixel size |
| $\Delta t$ | Time interval between two frames |
| $c(x, t)$ | Concentration field (function of space and time) |
| $I_k^{DATA}$ | Raw data vector |
| $I_k^{TH}$ | Noiseless theoretical data vector |
| $Y_i^{DATA}$ | Preprocessed (compressed) data vector |
| $Y_i^{TH}$ | Noiseless theoretical preprocessed data vector |
| $S$ | Sampling operator, maps concentration field onto raw data vector |
| $C$ | Compression operator, maps raw data vector onto preprocessed vector |
| $F$ | Spatial Fourier transform matrix |
| $n_q$ | Number of spatial modes kept per axis after compression |

Considering a diffusion model, we found an average value of the diffusion coefficient, ⟨*D*^*est*^ ⟩ = 1.93 10^−4^ *µm*^2^*s*^−1^, lower than diffusion coefficients of proteins in cell membranes (2), but in agreement with the order of magnitude − 10^−4^ 10^−2^ *µm*^2^*s*^−1^ for membrane-localised proteins in budding yeast or in fission yeast (8, 46–49). Such low value of surface diffusion coefficient is likely due to the presence of a cell wall (50). We also note that the empirical error —i.e. the standard deviation of the diffusion coefficient over all experiments — Δ*D*_emp_ = 1.40 10^−4^ *µm*^2^*s*^−1^, is much greater than the estimated error in single experiments⟨ Δ*D*_est_ ⟩ = 0.07 10^−4^ *µm*^2^*s*^−1^. This reflects biological cell-to-cell variability in diffusion, which has already been observed in cultured animal cells based on single particle tracking (51).

## 3 CONCLUSION

We have developed HiFRAP, a method to infer reaction (or exchange)-diffusion kinetic parameters from FRAP with imperfect conditions, and we have implemented it as a Fiji plugin. HiFRAP uses a whole time-lapse sequence and the variable projection approach to derive kinetic parameters, errors on these parameters, and a test of model validity so as to select which model better explains experimental observations.

HiFRAP combines all useful features that have been developed separately in previous work. HiFRAP makes it possible to infer several parameters from a single FRAP experiment (9, 15, 33–36, 38–43), as long as system dynamics is sufficiently sensitive to these parameters (37). We make no assumption on the bleaching profile (17–20, 23, 40, 42) nor on boundary conditions; HiFRAP is thus well-suited to experimental conditions in which it is difficult to bleach uniformly the target region or to be far from boundaries. HiFRAP accounts for intrinsic photobleaching associated with repeated imaging, either based on a control region or on inference of photobleaching rate (19, 22, 40). Finally, HiFRAP integrates three features that do not seem to have been implemented before: errors on parameter estimation for a single experiment; diffraction in the microscopy setup — though the precise characteristics of the point-spread function are not required as long as its width is larger than pixel size; test of model goodness. We ran HiFRAP against classical benchmarks and found that HiFRAP provides lower or equal errors despite being more general.

However, HiFRAP has a few limitations. First of all, HiFRAP is relatively slow (execution time of a few seconds with standard parameters and standard computer). Most available methods do not explicitly deal with noise. We were led to make the simplifying assumption that noise is homogeneous and stationary, which is not necessarily true (40, 52). For example, the amplitude of the shot noise associated with photon counting is proportional to the square root of the signal. Such spatiotemporal variation of noise would likely affect error estimation but not parameter estimation.

In principle HiFRAP can be adapted to FRAP variants, involving for instance continuous photobleaching of a region or photoconversion of a fluorophore (5)), and to the FRAP analysis of other types of dynamics, such as advection by active transport (9, 43), sub-diffusion (26, 27), multiple-species (9, 37, 40), and non-flat geometries (24, 25). HiFRAP assumes linearity of underlying partial differential equations (PDEs), but this is not a strong limitations as the dynamics becomes quickly linear upon return of the system to its equilibrium state.

Our method is actually not restricted to FRAP and could be used for inference of parameters for any linear PDE based on the effects of a perturbation on the system. We can therefore expect applicability of our method to capillary isoelectric focusing (53) or to optogenetics (54). Overall, our approach can be considered as a good alternative to machine learning approaches since it does not require training (55).

## 4 THEORY AND METHODS

### 4.1 Data sampling and compression

Here and elsewhere, we present our methodology for a 2-dimensional system, which also applies to a 1-dimensional system unless specified. We consider a spatiotemporal signal collected at discrete positions *X* ^(1)^, *X* ^(2)^ such that *X* ^(1)^ = *X* ^(2)^ = [0, Δ*x*, …, (*n*_*x*_ −1) Δ*x*] and times *T* = [0, Δ*t*, …, (*n*_*t*_ −1) Δ*t*], where Δ*x* is the spatial mesh size and Δ*t* is the time step, *n*_*x*_ × *n*_*x*_ is the number of pixels and *n*_*t*_ the number time frames. We vectorise the measurements by arranging them into a unique vector 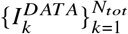 composed of 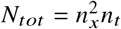 elements, where the index 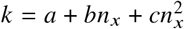 is associated to the space-time triplet 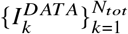 and *T*_*a*_ being the *a*-th components of the vectors *X* ^(*p*)^ and *T*, respectively. We assume that the empirical data correspond to a theoretical spatio-temporal signal *c*(*x, y, t*) that can be sampled into a theoretical data vector through the sampling operator 𝒮 :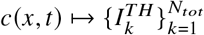, defined as 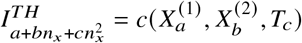.

To save data storage space and computational time, we compress these vectors into smaller vectors made of *N* elements (*N < N*_*tot*_), defining the compression operator 𝒞 : 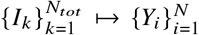. In practice, we chose to compress in the space domain because *n*_x_ × *n*_*x*_ is in general much greater than *n*_*t*_ : We keep *n*_*q*_ spatial Fourier coefficient per axis, so that 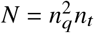 . Accordingly, we define the compression operator as

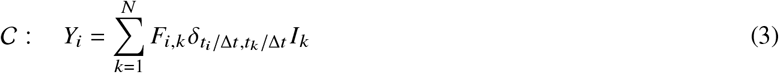

where *δ*_*i*,*k*_ is Kronecker’s delta and the Fourier matrix has elements 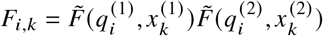 with

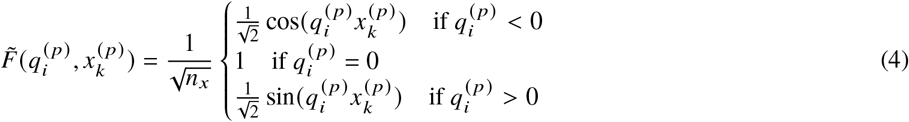

where the index *i* is associated to the Fourier-temporal vectorization with the Fourier vector of each axis spanning the value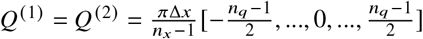

The results in Figure S2 show that as long as *n*_*q*_ ≥ 3, compression affects neither the accuracy — how close the mean of the distribution of the estimated parameter is to the true value — nor the precision of the estimation — the variance of the estimated parameter distribution. Typically, *n*_*q*_ ≥ 5 is sufficient to reach 80% of the precision that would be obtained without compression.

### 4.2 Inferring dynamical parameters

#### 4.2.1 Problem formulation

Our aim is to estimate the vector 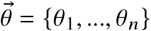 of the parameters of a linear PDE from an empirical signal 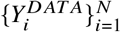. We consider system dynamics following a linear perturbation *w* (*x*). The theoretical, noiseless solution to the PDE takes the form

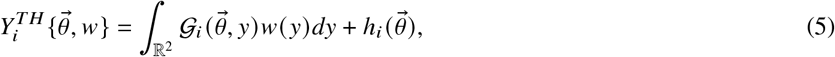

where the linear operator 𝒢 (related to Green’s function) and the shift vector *h* are specific to the PDE. As we will see later, it is also possible to include in 𝒢_*i*_ the effect of any linear operation on the signal, such as spatial filtering by the optical setup.

We consider experimental/technical noise, defined as 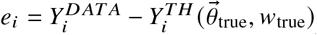, where *θ*_true_ and *w*_true_ are the true parameters and initial condition. We assume that the *e* _*i*_ are uncorrelated, i.e. ⟨*e*_*i*_ *e*_*j*_ ⟩ = *δ*_*i, j*_ *η*^2^, where *δ*_*i, j*_ is the Kronecker delta and *η* the unknown noise amplitude.

#### 4.2.2 Fitting the initial condition at fixed model parameters

Under these hypotheses, we can estimate the vector 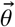 resorting to the least squares method, which consists in minimizing a cost that quantifies the differences between observed dataset and theoretical solution 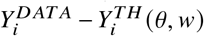. Since the initial profile *w* is not known, at any given *θ*, we first estimate it by minimising the cost 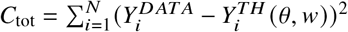, with respect to . By exploiting the fact that the theoretical solution 5 is linear in *w*, this operation can be performed analytically, yielding

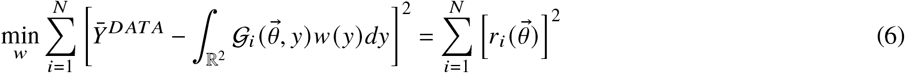

where the residual vector *r*, which represents the difference between the data vector and the constrained fit at fixed *θ*, is given by

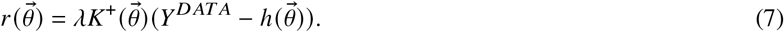

*K*^+^ is the pseudo-inverse of the kernel matrix *K* (56)

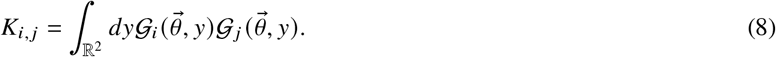

We use the pseudo-inverse *K*^+^,

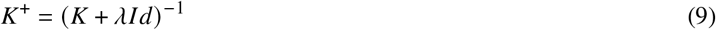

because the kernel matrix *K* is not definite positive. The small positive number *λ* ensures that the pseudo-inverse is well-defined. We take *λ* = *Nσ*_0_*ϵ*, using the largest eigenvalue *σ*_0_ of the kernel matrix *K* and machine precision *ϵ* (57), yielding typical values of *λ*(*θ*) in the range 10^−11^-10^−12^.

#### 4.2.3 Kernel reduction

To accelerate the computation of the pseudo-inverse in Eq. (7), we take advantage of the fact that multidimensional problems can often be reformulated in terms of smaller matrices that describe the structure of the kernel along each spatial direction. As we will see in Sections 4.4 for diffusion and 4.5 for reaction-diffusion, the kernel matrix can be factorized as

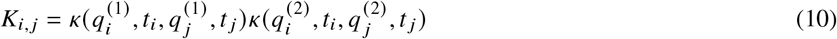

The values of the function *κ* at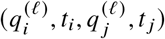, with *ℓ ∈* {1, 2}, are collected in a matrix *κ* ^(*ℓ*)^ of dimension *n*_*q*_*n*_*t*_ × *n*_*q*_*n*_*t*_, whose *u*-th row and *v*-th column are mapped by the relations *u* = *a* + *cn*_*q*_, *v* = *a*^′^ + *c*^′^*n*_*q*_ for *κ* ^(1)^ and *u* = *b* + *cn*_*q*_, *v* = *b*^′^ + *c*^′^*n*_*q*_ for *κ* ^(2)^, where *a, b, c* are associated to the triplet 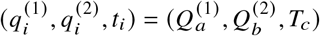 and *a*^′^, *b*^′^, *c*^′^ to 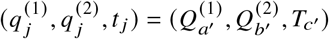 as previously defined in section 4.1.

Using singular value decomposition (SVD) based truncation (58), we approximate the matrices *κ* ^(*ℓ*)^ by considering the *n*_*K*_ eigenvalues *σ*_*k*_, *k ∈* {0, .., *n*_*K*_ − 1}, larger the same threshold *λ* = *n*_*q*_*n*_*t*_ *ϵ σ*_0_ as for the pseudo-inverse and the associated eigenvectors 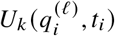. This operation yields an approximated kernel

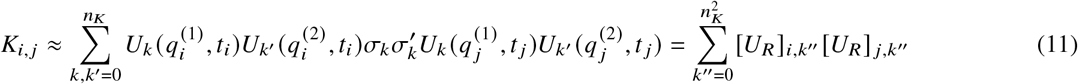

where *U*_*R*_ is a matrix of size 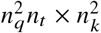, with components 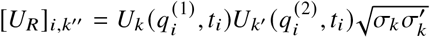 with *k*^′′^ = *k* + *n*_*k*_ *k*^′^. Under this approximation, exploing the Woodbury matrix identity, the pseudo-inverse takes the form

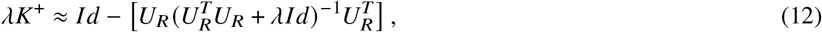

which is much faster to compute compared to its full expression since the size of the matrix 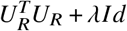 is smaller than the size of *K* + *λI d* by a factor of ∼10 with the parameters used here. We note that, because of the definition of *λ*, the errors induced by the approximations above are of the order of machine precision and so are negligible. In practice, we use this approximation of the pseudo-inverse *K*^+^ for a 2-dimensional system, but not in dimension 1 because the computational gain is limited, in which case we directly compute the pseudo-inverse of the kernel matrix as given by Equation (9).

#### 4.2.4 Fitting model parameters

The next step would be to minimise the sum of the squared residuals (Eq. 7) with respect to 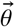 . However, the average contribution of the noise vector 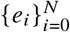 to the residuals 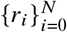 scales with the effective degrees of freedom 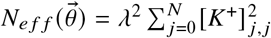 associated with the fit of the initial condition (59) whose value varies according to 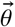. For this reason, to avoid biasing the inference towards values of 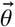 which requires stronger constraints for the initial condition fit (indicated by a lower number of effective degrees of freedom), we normalize the sum of residuals squared in Equation 7 by 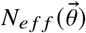. Combining this normalisation with Equations (7 and 9), we obtain the estimation of the parameter vector by minimizing the cost 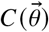 as

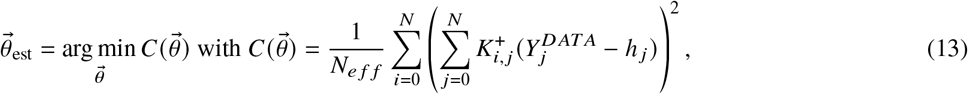

where *N*_*e f f*_, *K*^+^, *h* and *λ* all depend on 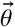.

For a 2 dimensional system, the kernel reduction 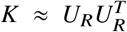 is used to compute both the pseudoinverse, based on Equation (12), and the effective degrees of freedom at the denominator of Equation (13), based on 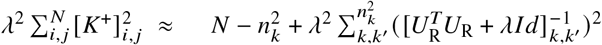 . For a 1 dimensional system, the cost in (13) is directly computed with the primary definition of the kernel (Eq. 8).

#### 4.2.5 Error on parameter estimation and goodness of the fit

Once the parameters 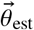 have been inferred by minimizing the cost function (13), the error on their estimation can be calculated using a quadratic expansion of the cost function around its minimum, yielding an estimated error matrix

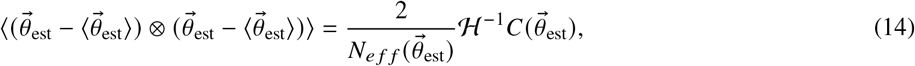

where ℋ is the Hessian of the cost function with respect to 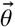 evaluated at 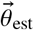.

We use two approaches to assess the goodness of the fit. The first is based on the adjusted coefficient of determination, which is the coefficient of determination (usually denoted by *R*^2^) corrected using the number of degrees of freedom — this avoids a spurious increase of the coefficient of determination with the number of fitting parameters (60). In our case, the adjusted 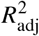 takes the form

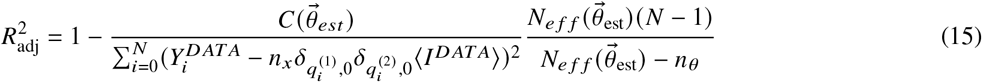

where 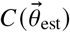 is the minimum of the cost function (Eq. 13), ⟨*I* ^*DAT A*^⟩ is the average of the raw data vector and *n*_*θ*_ the length of the vector 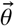. Larger (and closer to 1) *R*^2^ means a better fit (as this correspond to smaller minimum of the cost function). This test can be used to compare different models, e.g diffusion and reaction, for the same dataset *Y* ^*DAT A*^; the model with larger *R*^2^ should be favored. However, the adjusted *R*^2^ cannot be used for an absolute evaluation of the goodness of the fit because the standard deviation of the data vector may vary from experiment to experiment, in particular with the shape and depth of the initial bleaching profile.

For an absolute evaluation of the goodness of the fit, we use a Kolmogorv-Smirnov statistical test as follows. The starting point is that multiplication by *λK*^+^ in Eq. (7) approximately corresponds to projection on the orthogonal space of the matrix *K* (or of *U*_*R*_ in the case of kernel reduction). As consequence, the residuals can be approximated as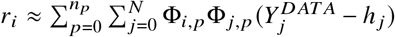, where the columns of the null matrix F (size *n* _*p*_ × *N*) are the eigenvectors of *K* (or le ft eigenvectors of *U*_*R*_) associated with eigenvalues above the threshold *λ* = *Nσ*_0_*ϵ* . If we define the projected residual as 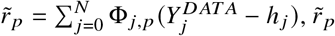 computed at 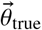depends only on the no is e contribution 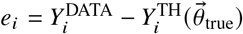 because 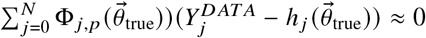 . Therefore, since 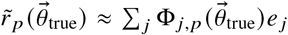 is a large linear combination of uncorrelated noise, the projected residuals computed at the true kinetic parameter values should be normally distributed. Accordingly, we apply the Kolmogorov-Smirnov normality test (45) to the vector *r*. We test the hypothesis that the model is compatible with the data by testing the null hypothesis that the reduced residuals are normally distributed. As can be seen in Figure S5, we obtain a *p*_*value*_ smaller than 0.05 only for 5% of the simulations, as expected when the artificial data correspond to the model tested. However, this method is much slower compared to the adjusted *R*^2^, as the computation of eigenvectors scales with O(*N*^3^), while the computation of the standard deviation scales with O(*N*).

### 4.3 Modelling signal acquisition

Here we aim at accounting for two experimental features: photobleaching during imaging associated with imaging and diffraction in the optical setup. When the sample is imaged, photobleaching occurs at a rate *ε* per image. Diffraction implies that the detectors collect the true signal convolved by the point-spread function, assumed to be a Gaussian of width *µ* as a point-spread function. The sampling operator can then be expressed as

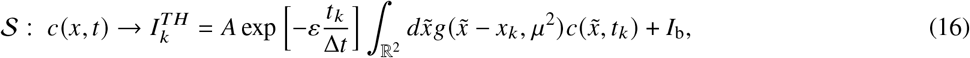

where the scaling parameter *A* can be set to 1 if the unit of intensity is arbitrary, the two-dimensional point-spread function is given by

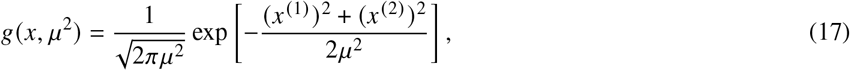

and *I*_b_ is the background signal.

### 4.4 Inference in the case of pure diffusion without photobleaching

Here we apply the inference method to the signal 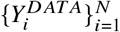 in the case of a purely diffusive model when photobleaching during imaging is negligible, i.e. *ε* = 0. In this case, the only kinetic parameter to infer is *D*, the diffusion coefficient. The analytical solution of diffusion PDE 1 with a generic initial condition *c*(*y*, 0) can be expressed as

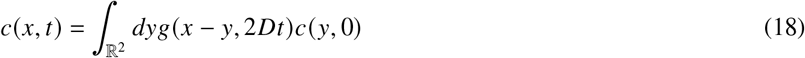

with *g* defined as in Equation 17. If we apply the sampling operator S to *c*(*x, t*) and the compress the resulting vector 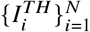, we obtain the theoretical compressed signal as

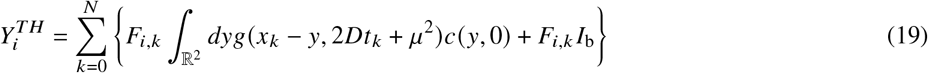

By comparison with Equation (5), we identify the kernel matrix 𝒢_*i*_ (*x, y*) = ∑_*k*_ *g* (*x*_*k*_ −*y*, 2*Dt*_*k*_ +*µ*^2^), the shift vector *h*_*i*_ = 0 and *w* (*y*) = *c* (*y*, 0) + *I*_*B*_. We remark that in this case there is no need to estimate the background value to infer diffusion, as its contribution is incorporated into the initial perturbation *w* (*y*) . Moreover, even if the width of the PSF *µ* is not known, in practice we can set this parameter equal to Δ*x*, the pixel size. Indeed, we found that the average and the variance of the estimated parameter distribution are not affected by this choice as long as the true value of the system *µ*_true_ *≳* 1, see Figure S4. We obtain the kernel matrix from Equation (8) as

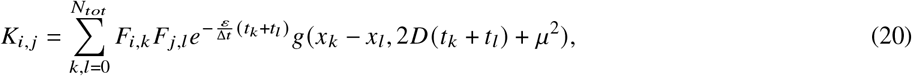

from which it is possible to compute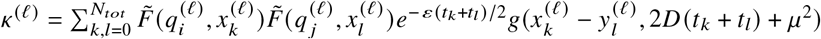, with *ℓ ∈* {1, 2}, and *U*_*R*_ with Eq. (11). Finally, we infer the diffusion coefficient *D*_est_ by minimising the cost function (Eq. 13).

### 4.5 Inference in the case of reaction-diffusion with photobleaching

Now we consider the underlying model to be a linear reaction-diffusion PDE (Equation 2) and photobleaching during imaging not to be negligible, i.e. *ε* ≠ 0. Following the same approach as in the previous section, the theoretical compressed signal takes the form

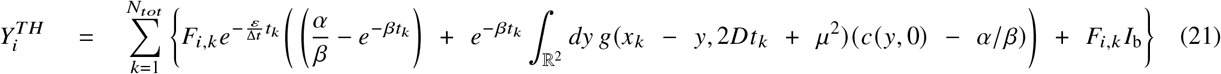

By comparison with Equation (5), we identify 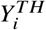 and we obtain the kernel from Equation (8)

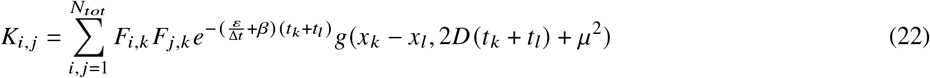

and

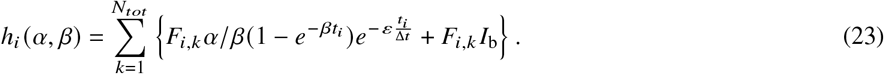

We not e that *h*_*i*_ (*α, β*) is defined at *β* = 0 by continuity using lim_*β*→0_(1 − exp(−*βt*_*i*_))/*β* = *t*_*i*_. From Eq. (22), we deduce 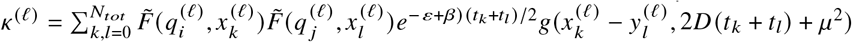, with *ℓ ∈* {1, 2}, and *U*_*R*_ with Eq. (11).

Finally, the cost function defined in Equation (13) can be minimised with respect to the kinetic parameters *D, α, β* and *I*_BG_. Since *I*_BG_ and *α* are linear parameters, the values, 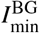 and *α*_min_, that minimise the cost for fixed *D* and *β* can be obtained analytically as

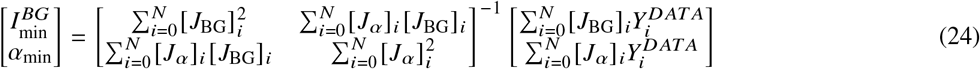

where 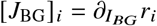 and [*J*_*α*_]_*i*_ = *∂*_*α*_*r*_*i*_ are the jacobian of the residuals 7 with respect to *I*_BG_ and *α* taking the form

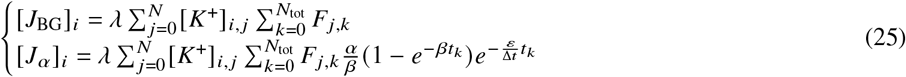

So, the cost in equation (13) is computed directly at 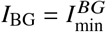 and *α* = *α*_min_ and minimised numerically with respect to *D β*, and *ε*, yielding the final estimations 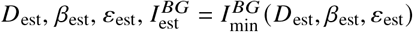 and *α*_est_ = *α*_min_ (*D*_est_, *β*_est_, *ε*_est_).

### 4.6 Improving parameters estimation using extra-information

Even though we can estimate all parameters in Eq. 21, knowing *a priori* the background and the photobleaching decay rate can improve the estimation of model parameters *D*_est_, *α*_est_ and *β*_est_. For this reason, it is important to take advantage of information available beyond the FRAPped area. A sample-free area gives access to the background *I*_BG_ by averaging the signal over this area. The photobleaching loss-rate can be extracted from a region of the sample far enough from the FRAPped area, by an exponential fit 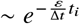 to the spatial average signal over this area. Moreover, if a pre-FRAP image is present, we can obtain the stationary concentration *c*_*s*_ by averaging the signal over the area that is later FRAPped. This value sets the ratio between the exchange and source rate, *c*_*s*_ = *α* /*β*. This *a priori* information can be integrated in the computation of the kernel and shift terms. In table 5, we recapitulate the kernel and shift terms for all experimental conditions: with or without photobleaching, with or without an unfrapped control area, with or without a sample-free area and with or without prebleaching images. It is important to remark that when photobleaching is absent and no background area is available, the contribution to the signal intensity of the source rate *α* and the background value *I*_BG_ cannot be distinguished. In such cases, HiFRAP can only estimate the combined parameter *α*^∗^ = *α* + *βI*_BG_. To determine *α* separately, the user must obtain the value of *I*_BG_ from independent experiments.

**Table 3:** Notations for inference.

|  |  |
| --- | --- |
| $\vec{\theta}$ | Vector of kinetic parameters to be inferred |
| $\vec{\theta}_{true}$ | Vector of true values of kinetic parameters |
| $\vec{\theta}_{est}$ | Vector of estimated kinetic parameters |
| $w$ | Spatial perturbation |
| $\mathcal{G}w + h$ | The linear operator $\mathcal{G}$ and the shift vector $h$ |
| $e_i$ | Noisy part of the preprocessed vector |
| $\eta$ | Amplitude of the noise |
| $K$ | Kernel operator |
| $K^+$ | Pseudo-inverse of the kernel operator |
| $\lambda$ | Small positive number to ensure matrix inversion or threshold on eigenvalues |
| $U_R U_R^T$ | Reduced kernel by SVD truncation along each axis |
| $N_{eff}$ | Trace of $[\lambda K^+]^2$ representing the effective degrees of freedom |
| $\Phi$ | Null matrix associated with kernel operator |
| $r$ | Residual between data and theory vector after fitting the initial condition at $\vec{\theta}$ fixed |
| $\tilde{r}$ | Projected residual in the null space generated by $\Phi$ |

**Table 4:** Parameters of signal acquisition model and of PDE.

|  |  |
| --- | --- |
| $I_{BG}$ | Background signal |
| $\mu$ | Width of point-spread function |
| $\varepsilon$ | Rate of photobleaching during imaging |
| $D$ | Diffusion coefficient |
| $\alpha$ | Source rate |
| $\beta$ | Exchange (or dissociation) rate |

**Table 5:**
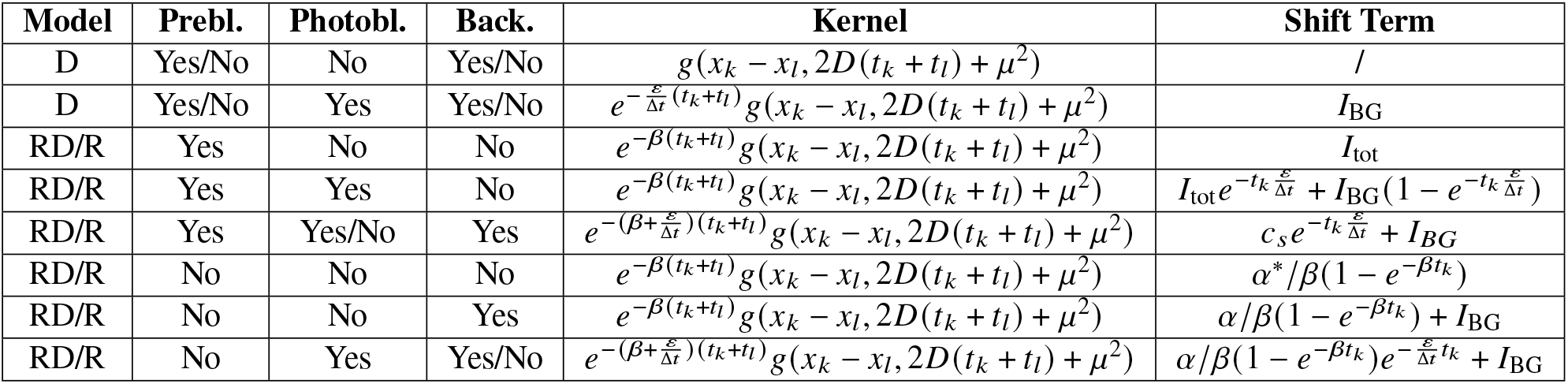
Summary of the kernel and shift term formulas for each model under different experimental conditions. From left to right: model (diffusion — D, reaction — R, or reaction-diffusion — RD), presence of prebleaching images (yes or no), whether imaging induces photobleaching (yes or no), whether a sample-free background area is available (yes or no), and corresponding kernel and shift term (without compression for ease of notation). The diffusion coefficient, *D*, and exchange rate, *β*, are always estimated by numerical minimization of the cost function or set to zero in the formulas when not included in the model. The background signal, *I*_BG_, is estimated by averaging the signal in the background area when this information is available; otherwise, if *I*_*B*_ appears in the formula, its estimated value is determined through analytical minimization of the cost function. The stationary concentration (in arbitrary units), *c*_*s*_, is estimated by averaging the prebleaching images and subtracting the background *I*_BG_ when both prebleaching and background areas exist. When the background does not exist, the average prebleaching signal corresponds to *I*_TOT_, the sum of the background signal and the stationary concentration. The source rate, *α*, is determined via analytical minimization of the cost function or a posteriori as *α*_est_ = *c*_*s*_/ *β*_est_ when *c*_*s*_ is known, or as *α*_est_ = (*I*_tot_ − *I*_BG_) /*β*_est_ when the background is estimated through cost minimisation. In the other cases (third and sixth row), it is not possible to estimate directly *α* but only the parameter *α*^∗^ = *α β*+*I*_*BG*_ from analytical minimisation of the cost function (sixth row) or a posteriori from 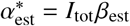 if *I*_tot_ is known (third row). The photobleaching decay rate *ε* is either computed from the control area when present or by numerical minimisation of the cost function. Δ*t* is the time interval between two images while the function *g* is defined in Eq. 17.

### 4.7 Numerical implementation

In this section, we describe the numerical implementation of HiFRAP, using `SciPy` and `NumPy` package in Python. We wrapped the corresponding scripts in an ImageJ plugin also called HiFRAP.

We begin with the calculation of the cost function, which requires computing the inverse of a matrix (*K* +*λI d*) ^−1^ (or its expression with *U*_*R*_ in Eq. 12). This is achieved using the Cholesky decomposition. Specifically, we first compute the Cholesky factor *L* using `scipy.linalg.lapack.dpotrf`. The inverse, *L*^−1^, is then obtained using `scipy.linalg.lapack.dtrtri`, which takes advantage of the triangular shape of *L* using a recursive block algorithm. Finally the pseudo-inverse is given by (*K* +*λI d*) ^−1^ = *L*^−*T*^ *L*^−1^. To obtain the threshold *λ*, the first eigenvalue is calculated by Lanczos Bidiagonalization using `scipy.linalg.svds.`

To minimize the cost function (Eq. 13) with respect to *D, β*, and/or *ε*, we first determine an appropriate initial condition for these parameters to start from the numeric algorithm. In this context, we suppose the typical relaxation time of diffusion 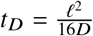, dissociation rate *t*_*β*_ = log(2)/*β* and/or photobleaching *t* _*ε*_ = Δ*t* log(2)/*ε* to be equal to half of the total observational window *T* . Since the typical bleaching size *ℓ* is not known, we approximate this value *ℓ ≈L* /2, where *L* is the total spatial window. These yields as initial parameters *D*_in_ = (*L* /2) ^2^ /16 /*T* (or (*L*/ 2) ^2^ /8/ *T* in 1D), *β*_*in*_ = log (2) /*T* and/or *ε*_*in*_ = log (2) Δ*t* /*T*_tot_

Next, the full reflective trust-region algorithm is applied as implemented in `scipy.optimize.least_squares(method=‘trf’`). Optimization is performed by restricting the parameter ranges according to the spatiotemporal observation window, as parameter values outside these ranges would be physically meaningless. For the diffusion coefficient *D*, we impose that the timescale associated with the slowest Fourier mode, of order *L*^2^ /*D* (where *L* denotes the total spatial window), must be larger than 5Δ*t*, ensuring sufficient temporal resolution. Conversely, the timescale associated with the fastest Fourier mode, of order (*L* /*n*_*q*_) ^2^ /*D* (with *n*_*q*_ the highest wavenumber considered), must be smaller than 40*T*, where *T* denotes the total temporal observation time. For the exchange rate *β*, whose characteristic timescale is of order 1/ *β*, we constrain *β* such that −*T <* 1 /*β <* 100*T* . Here, negative values of 1 /*β* are allowed, as they can arise when the empirical error Δ*β*_emp_ is comparable to the true value *β*_true_. For the photobleaching decay rate *ε*, we require that it is not negligible over the temporal window, restricting its values such that 100*T <* 1/*ε < T* /15. During the optimization procedure, convergence is considered achieved when the relative variation in the estimated parameter vector 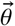 between successive iterations satisfies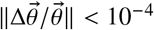.

Once the minimum of the cost function is found, the Hessian is computed using the Jacobian of residuals 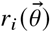 divided by square root of the trace 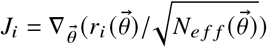, using the approximation 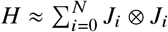, as implemented in the reflective trust-region numerical method. The Jacobian is computed numerically by taking a finite step of the order of 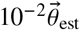, which corresponds to the typical noise error, except that the derivatives with respect to the source rate are computed analytically from Eq. 25. Finally, to evaluate the goodness of the fit, the null matrix Φ is obtained by computing the eigenvectors and eigenvalues of *K* or *U*_*R*_ using the singular value decomposition as implemented in `numpy.linalg.svd.`

### 4.8 Artificial data

Artificial concentration fields *c* (*x, t*) were obtained by solving analytically the reaction-diffusion equation (2) (or the diffusion equation 1), with known parameters *D*_*true*_, *α*_*true*_ and *β*_*true*_. The initial condition describes a square FRAPped profile,

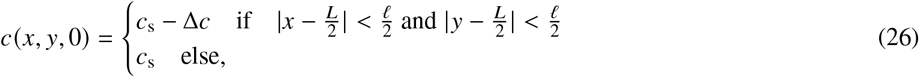

where Δ*c*/ *c*_s_ is the proportion of bleached fluorophores and *ℓ* is the square side length. To get the signal vector, the concentration solution was multiplied by a factor *A* and convoluted with a Gaussian function of width *µ*_true_ to mimic the effect of the PSF. Photobleaching during imaging is readily accounted for by multiplying the solution by an exponentially decaying function with rate *ε*_true_. The background value 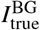 was added to the resulting signal. The theoretical data vector takes the form

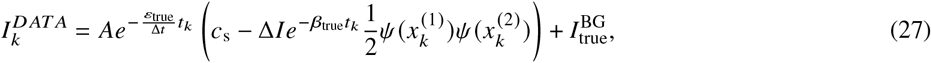

with

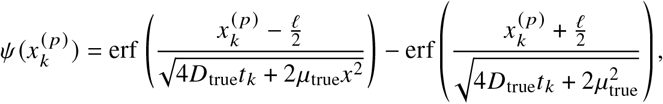

where the error function is defined as 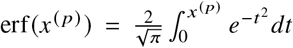 and Δ*I* = *A*Δ*c* the signal drop-off. The stationary concentration *c*_s_ is fixed for pure diffusion, whereas for reaction-diffusion, *c*_s_ = *α*_true_ /*β*_true_. The same procedure was applied for a Gaussian bleaching profile, X-shape and E-shape (obtained by translation, rotation, extension and superposition of of square bleaching profile)

To obtain a realistic dataset we add noise to the deterministic solution,

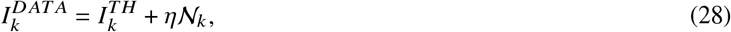

where *η* is noise amplitude and 𝒩_*k*_ is sampled from a Gaussian random variable of mean zero and standard deviation 1. Finally, we compress the simulated vector (Eq. 27) using the compression operator C (Equation 3).

Unless specified otherwise, we used the following default values: *n*_*x*_ = 121, *n*_*t*_ = 16, *ℓ*/(*n*_*x*_Δ*x*) = 3, *µ*_true_/Δ*x* = 1, *η*/Δ*I* = 0.25, *I*_BG_/Δ*I* = 0.5, *c*_s_/Δ*I* = 2 with Δ*x* = 1, Δ*t* = 1, Δ*I, A* = 1 in arbitrary units; for diffusive systems *D*_true_ = 2*ℓ*^2^/16/(*n*_*t*_ − 1)/Δ*t* or *D*_true_ = 7*ℓ*^2^/16/Δ*t*/(*n*_*t*_ − 1) (if photobleaching is present) and *β*_true_ = 0; for reaction-diffusive systems *D*_true_ = 7/2*ℓ*^2^/16/(*n*_*t*_ − 1)/Δ*t, α*_true_ = *β*_true_*c*_*s*_, *β*_true_ = 7/2 log(2)/(*n*_*t*_ − 1)/Δ*t*; if photobleaching is present *ε*_true_ = 1/(*n*_*t*_ − 1) otherwise *ε*_true_ = 0.

### 4.9 Experiments

The *Schizosaccharomyces pombe* strain *mtl2-GFP: ura4*^*+*^ (identifier RN21, (8)) was used for experimental validation of the method. Standard fission yeast methods and media were used (Forsburg & Rhind, 2006). The cells were grown in YE5S liquid culture overnight at 25^°^C, diluted in fresh medium and grown to an optical density (OD_600_) between 0.4 and 0.6 before live-imaging. Cells were imaged on EMM (minimal media) 2%-agarose pads at room temperature (22-25^°^C); EMM shows reduced background noise in comparison with YE5S agarose pads. Cells were imaged at their bottom surface, close to the coverslip. Images were acquired with a 100 × oil-immersion objective (CFI Plan Apo DM 100× /1.4 NA, Nikon) on an inverted spinning-disk confocal microscope equipped with a motorized stage and an automatic focus (Ti-Eclipse, Nikon, Japan), a Yokogawa CSUX1FW spinning unit, a Prime BSI camera (Teledyne Photometrics,USA) and an iLas2 module (GATACA Systems, France) for FRAP. During FRAP, a 0.4 *µ m* square ROI was bleached with the 491nm laser at 20-60 % power and 30 repetitions and the fluorescence recovery was monitored for a time interval ranging from 7 − 10 *s* .

An online supplement to this article can be found by visiting BJ Online at http://www.biophysj.org.

## AUTHOR CONTRIBUTIONS

EL designed and implemented the theory. CMD performed experiments. NM supervised experiments. AB and AF designed and supervised the study.

## ACKNOWLEDGMENTS

This work was funded by Agence Nationale de la Recherche (project CellWallSense, ANR-20-CE130003-01, to NM and AB).

## 5 SUPPLEMENTARY FIGURES

**Figure S1:**
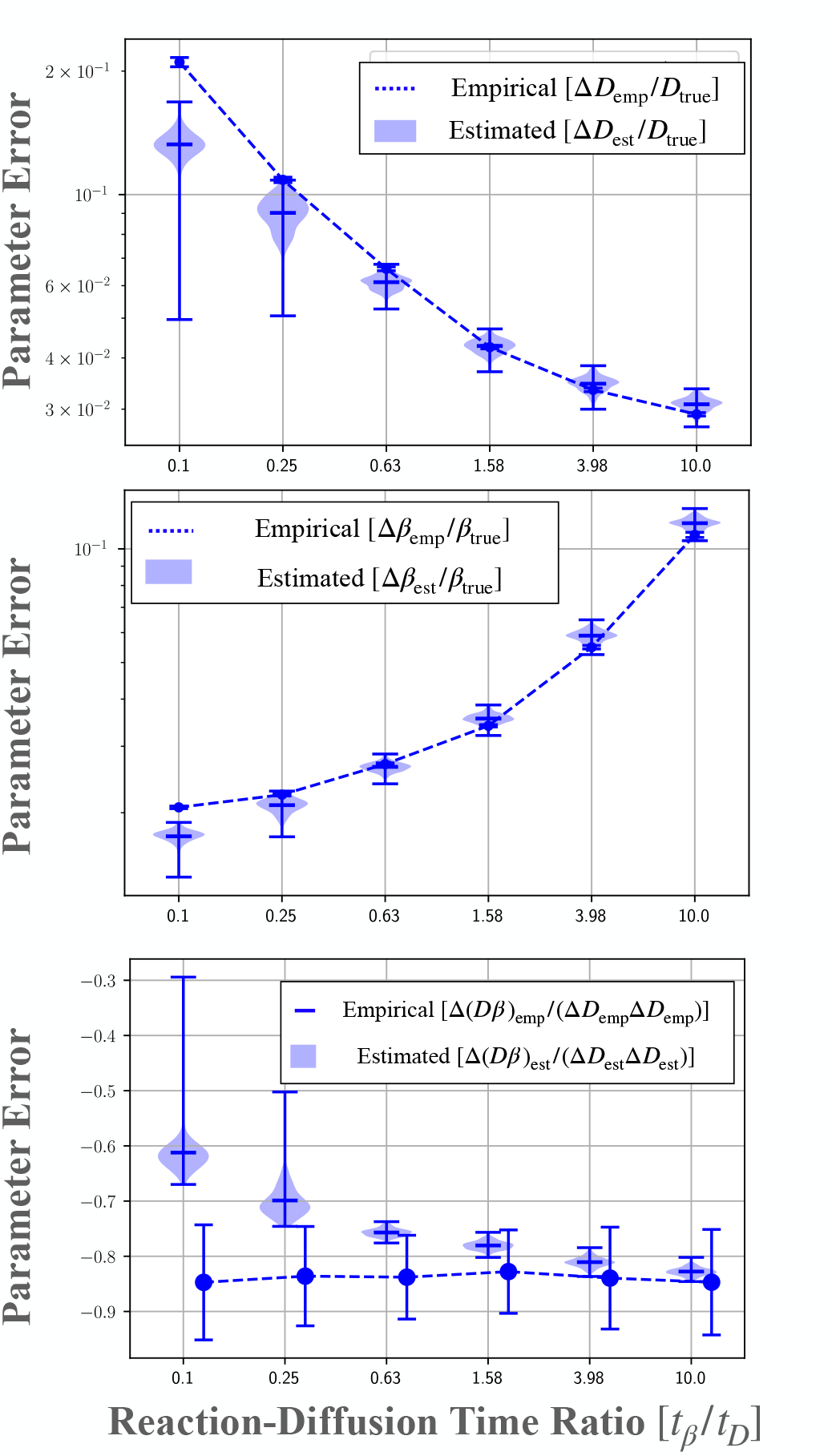
Comparison between empirical error (measured over all realisations of noise) and estimated error (measured from single realisations) when inferring diffusion coefficient *D* and exchange coefficient *β* as function of the reaction-diffusion time ratio *t*_*β*_/*t*_*D*_ = log(2)/*β*_true_16*D*_true_/*ℓ*^2^. From top to bottom: error on *D*, error on *β*, and correlation between *D* and *β*.

**Figure S2:**
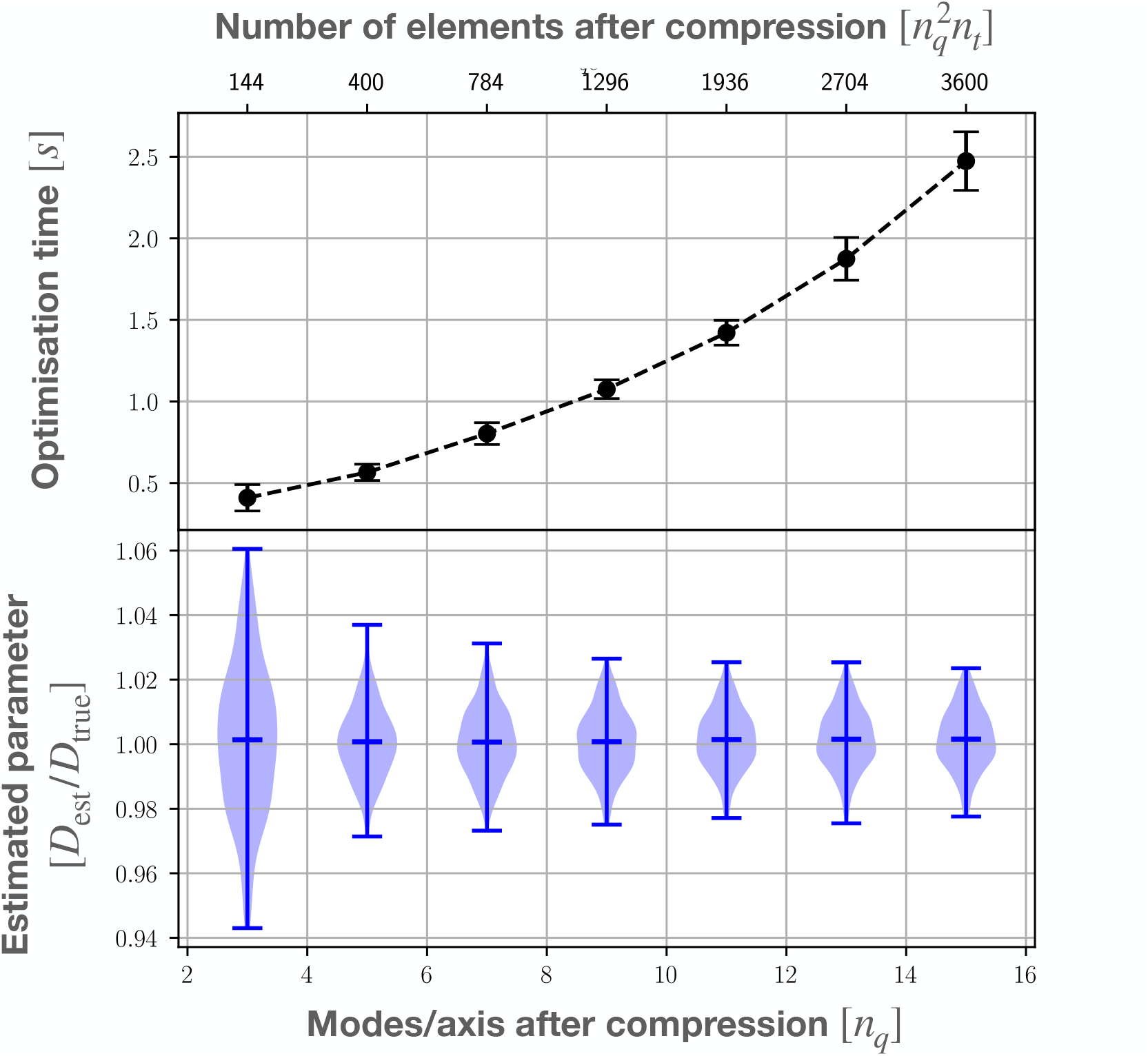
Effect of compression. Top: time necessary to numerically minimise the cost function *C* (*D*), where *D* is the diffusion coefficient, plotted as a function of the number of elements kept after compression 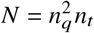, with *n*_*q*_ the number of modes per axis. Results were obtained only from one execution. Bottom: violin plot for 200 simulations showing the distribution of estimated coefficient of diffusion *D*_*est*_ (normalised with its true value *D*_*true*_) as function of the number of modes kept per axis after compression, *n*_*q*_

**Figure S3:**
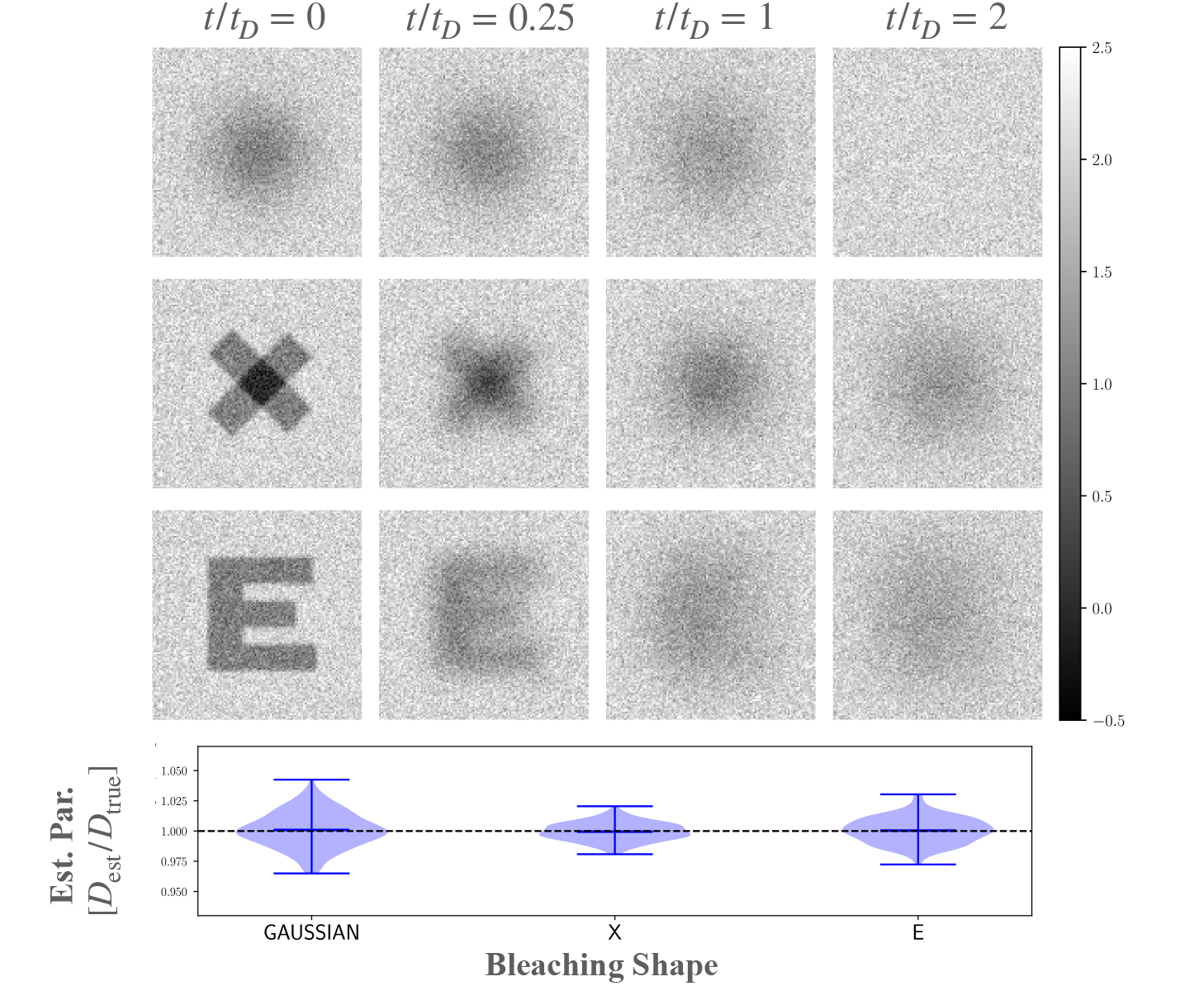
Top: Simulation of different diffusive dynamics with different bleaching shapes. The results are shown at different time (normalised by 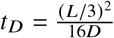, where *L* is the observation window size). The grayscale (left) corresponds to signal intensity normalised with the signal drop-off Δ*I*. Bottom: Violin plot of the distribution of estimated diffusion coefficient *D*_*est*_ (normalised with its true value *D*_true_) for different bleaching profiles. The dashed gray line represents the reference value *D*_est_/*D*_true_ = 1.

**Figure S4:**
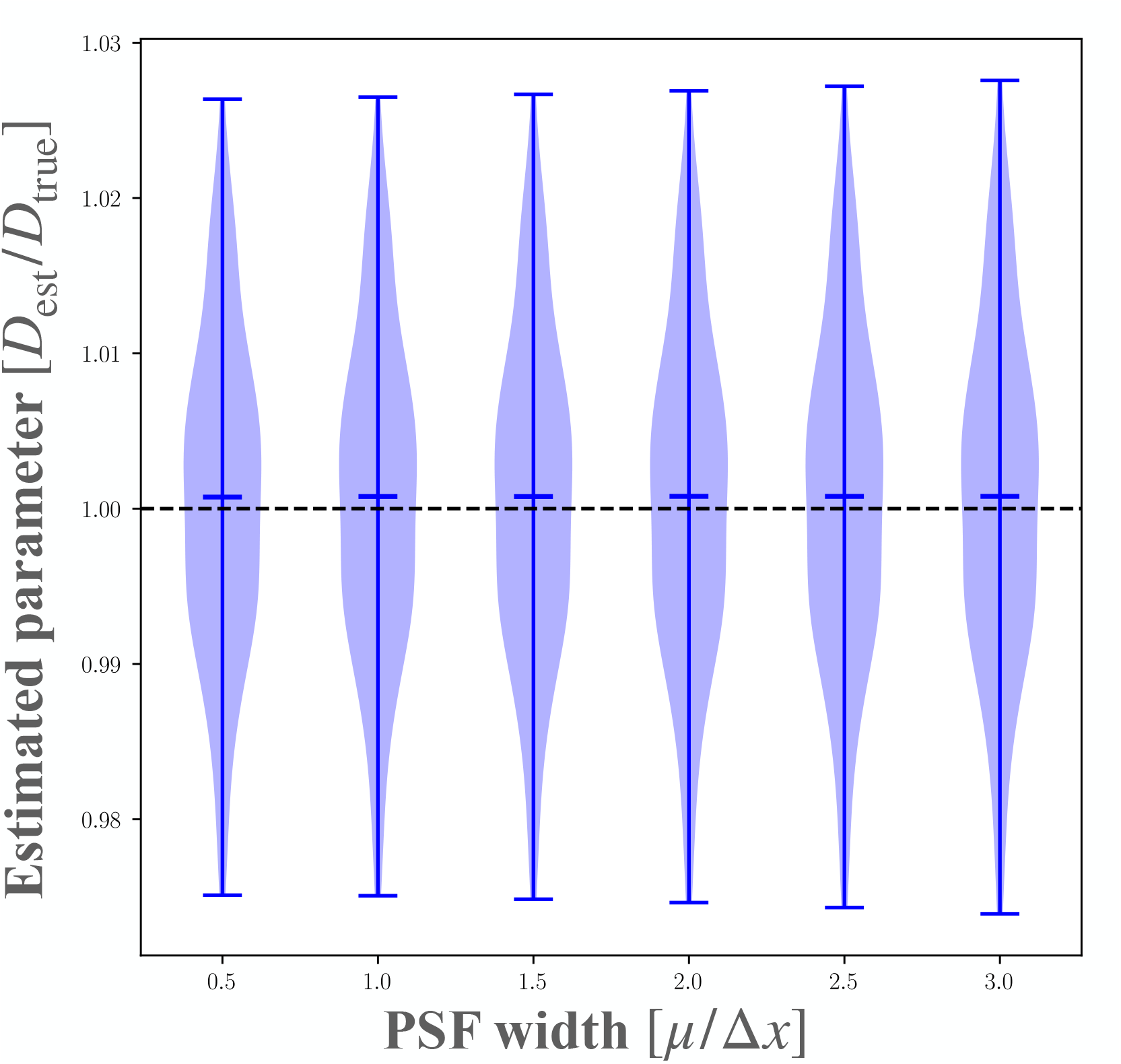
Effect of the point-spread function (PSF) width on estimation of the diffusion coefficient. Violin plot of the distribution of estimated coefficient *D*_*est*_ (normalised with its true value *D*_true_) for different PSF widths *µ* (normalised with pixel size Δ*x*). The dashed gray line represents the reference value *D*_est_/*D*_true_ = 1.

**Figure S5:**
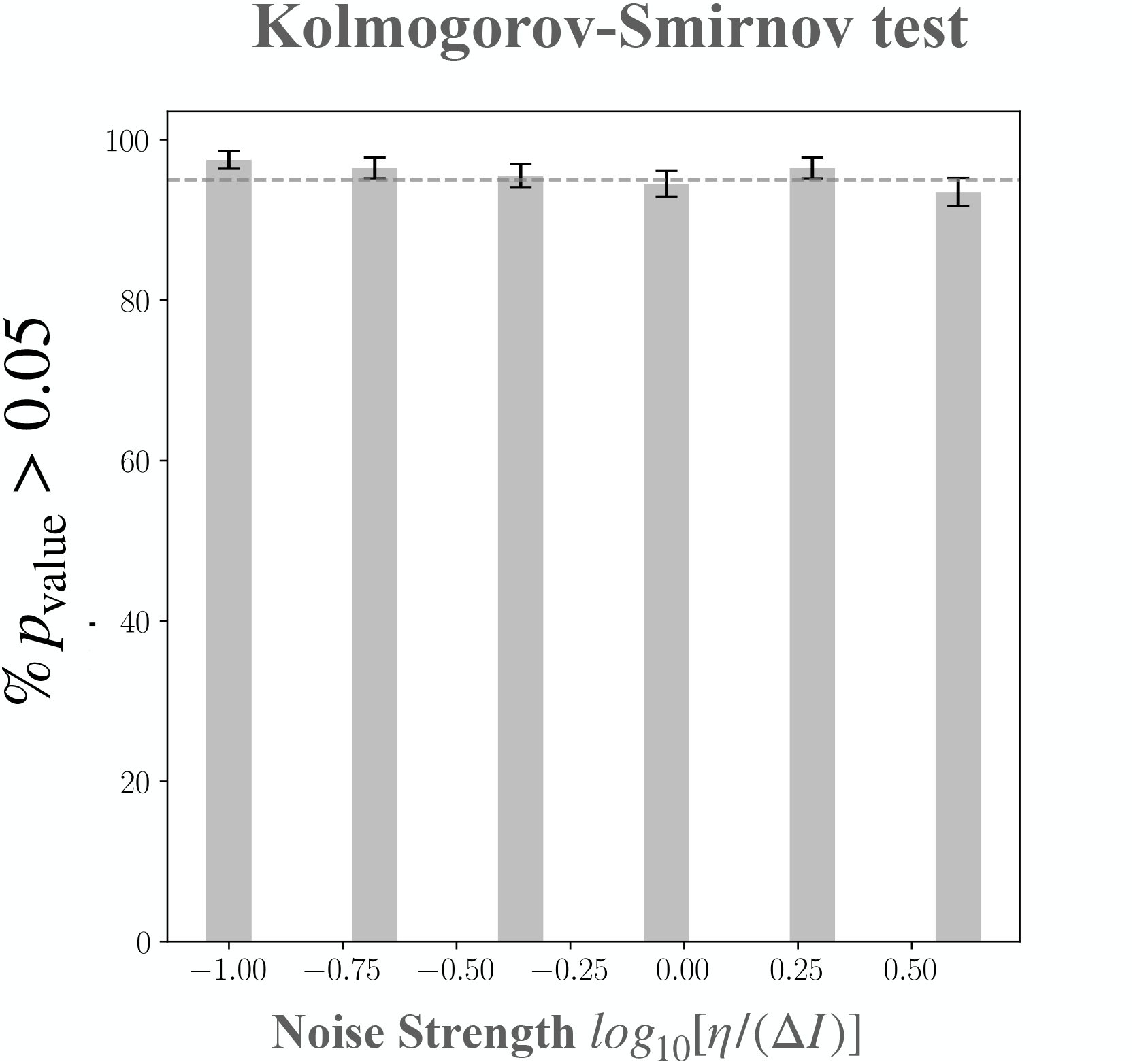
Kolmogorov test: the bar represents the percentage of times we obtain a Kolmogorov p-value greater than 0.05 after fitting data with the estimated diffusion value as a function of the noise strength (*η*, the amplitude of the noise and Δ*I*, the signal drop-off). The test is applied on 200 synthetic datasets of a diffusive system without photobleaching during imaging. Errorbars correspond to the error in sampling a binomial distribution.

**Figure S6:**
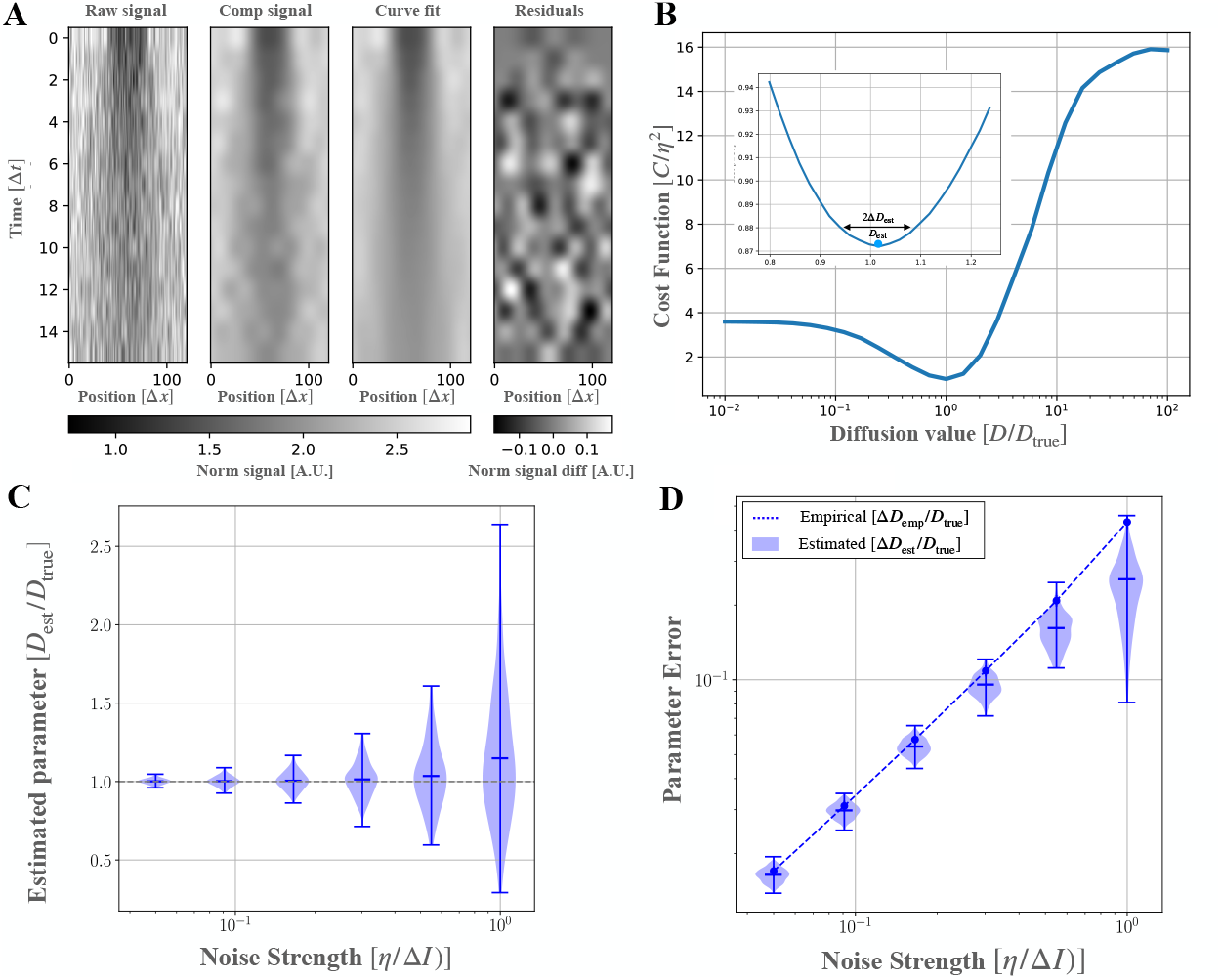
Inferring the diffusion coefficient from simulated FRAP in a 1D system. **A**. A line of length *L* is monitored and a central line region of side length *ℓ* = *L* /3 is FRAPped at *t* = 0. From left to right, kymographs (spatio-temporal representations) of: raw artificial data, microscope-like synthetic data accounting for diffraction and technical noise, compressed synthetic data, concentration field fitted by HiFRAP, and residuals of the fit. Grayscales indicate the concentration or signal intensity normalised by the drop in concentration Δ*I* at *t* = 0 in the FRAPped region. The decay rate per image due to photobleaching is set to *ε* = 0. For other parameters, default values are given in Section 4.8. **B**. Modified cost function *C* (normalised by noise amplitude *η*) as a function of fitting diffusion coefficient *D* (normalized by its true value *D*_true_), with a magnification of the neighbourhood of the minimum of *C* in the inset. The cost function is minimal at *D*_est_ = 1.01*D*_true_, which is close to *D*_true_, up to an estimated error Δ*D*_est_ = 0.07Δ*D*_true_. The total observation time *T* for this dataset is *T* = 2*t*_*D*_ and the time step Δ*t* = 0.133*t*_*D*_, with *t*_*D*_ = *ℓ*^2^ /8 /*D*_true_ the characteristic recovery time.**C-D** Validation on a collection of synthetic data. **C** Estimated diffusion coefficient *D*_est_; **D** estimated uncertainty Δ*D*_est_ and empirical error Δ*D*_emp_. The quantities are all normalised by the true diffusion constant *D*_true_ and plotted as a function of the normalised noise amplitude *η* /Δ*I*. Violins represent distributions of *D*_est_ and Δ*D*_est_ while the ticks highlight average and extreme values. The dashed gray line in **C** represents the reference value *D*_est_/*D*_true_ = 1, while the dashed blue line in **D** corresponds to the empirical error Δ*D*_emp_. The number of realisations is 200 for each value of noise strength.

